# Immunoregulatory subtype of dermal lymphatic endothelial cells at capillary terminals drives lymphatic malformations

**DOI:** 10.1101/2022.05.22.492950

**Authors:** Milena Petkova, Marle Kraft, Simon Stritt, Ines Martinez-Corral, Henrik Ortsäter, Ying Sun, Michael Vanlandewijck, Bojana Jakic, Eulàlia Baselga, Sandra D. Castillo, Mariona Graupera, Christer Betsholtz, Taija Mäkinen

## Abstract

Vascular malformations are congenital, chronically debilitating diseases. Somatic oncogenic mutations in *PIK3CA*, encoding p110α-PI3K, specifically cause venous and lymphatic malformations (LM), yet the basis of vessel type-restricted disease manifestation is unknown. Here we report endothelial subtype-specific responses to the common causative *Pik3ca^H1047R^* mutation, and reveal a new immunoregulatory subtype of dermal lymphatic capillary endothelial cells (iLECs) as a driver of LM pathology. Mouse model of *Pik3ca^H1047R^*-driven vascular malformations showed that cell proliferation was a common early response of venous and lymphatic ECs to oncogenic *Pik3ca*, but sustained selectively in LECs of advanced lesions. Lymphatic overgrowth was associated with increased pro-inflammatory cytokine levels and pro-lymphangiogenic myeloid cell infiltrate. Single-cell transcriptomics revealed a new LEC subtype at capillary terminals, characterized by the expression of immunoregulatory genes. Selective expansion and activation of iLECs in the *Pik3ca^H1047R^* mice was evidenced by proliferation and upregulation of pro-inflammatory genes. Importantly, macrophage depletion or anti-inflammatory COX-2 inhibition limited *Pik3ca^H1047R^*-driven lymphangiogenesis. This provides a therapeutic target for LM and suggests a paracrine crosstalk in which LEC-autonomous oncogenic *Pik3ca* signaling induces immune activation that in turn sustains pathological lymphangiogenesis. Identification of iLECs indicates that peripheral lymphatic vessels not only respond to inflammation but also actively orchestrate the immune response.

## Introduction

Venous malformations (VMs) and lymphatic malformations (LMs) are chronic diseases characterised by vascular lesions that range from simple skin discoloration to large deformations, or fluid-filled cysts to infiltrative soft-tissue masses, respectively (1, 2). They are often associated with significant morbidity, and in some cases life-threatening complications, due to pain, bleeding and functional impairment of nearby organs. Somatic activating *PIK3CA* mutations have been identified as causative of the majority of LMs (3, 4) and a smaller proportion of VMs (5–7). *PIK3CA* is frequently mutated also in cancer and other pathologies characterized by tissue hyperplasia, the so-called PIK3CA-related overgrowth spectrum (PROS) (8).

*PIK3CA* encodes the p110α subunit of the phosphoinositide 3-kinase (PI3K) that catalyzes the production of phosphatidylinositol (3,4,5)-triphosphate (PIP_3_) at the plasma membrane, leading to activation of downstream signaling cascades such as the AKT-mTOR pathway. PI3K signaling controls a variety of cellular processes in both blood and lymphatic vasculatures, including endothelial cell (EC) migration, survival and proliferation as well as vessel sprouting, thereby critically regulating vascular maintenance and growth (9, 10). Genetic loss-of-function studies in mice have uncovered a critical role of p110α in the normal development of blood and lymphatic vessels (11–13). Conversely, expression of an activating *PIK3CA* mutation in ECs led to vascular overgrowth and malformations in mice (14–16). EC-autonomous effects in the pathogenesis of both VM and LM are demonstrated by the presence of *PIK3CA* mutations specifically in ECs but not in other cell types (4, 17, 18).

*PIK3CA* mutations frequently occur in two hot spot regions encoding the helical domain and the kinase domain, with an H1047R substitution in the latter representing one of the most frequent VM/LM and cancer mutation (1, 2, 8). Identification of PIK3CA mutations as drivers of vascular malformations has enabled repurposing available FDA-approved inhibitors of the PI3K pathway for their treatment. For example, rapamycin that targets the PI3K-AKT downstream effector mTOR (mammalian target of rapamycin) has shown efficacy in relieving symptoms in VM and LM patients although it rarely results in the regression of lesions (1, 2). Apart from the identified EC-autonomous mutations driving vascular anomalies, emerging evidence points to synergistically acting paracrine mechanisms that contribute to disease progression (2). For example, increased paracrine vascular endothelial growth factor C (VEGF-C) signaling is observed in LMs in mice and human patients (16, 18, 19), and it is required for the growth of *Pik3ca*-driven LM in mice (16). Interestingly, inhibition of VEGF-C was more effective than rapamycin in limiting LM growth in mice, and when administered in combination with rapamycin it even promoted regression of the abnormal lymphatic vessels (16). Better understanding of both the aberrant EC-autonomous signaling and the paracrine mechanisms should aid the development of effective and targeted combinatorial therapies for LMs and other vascular malformations.

Here, we investigated the endothelial subtype-specific mechanisms underlying *PIK3CA*-driven LM in comparison to VM. Analyses of mouse models of *Pik3ca^H1047R^*-driven vascular malformations revealed lymphatic and blood vessel-type specific responses resulting in distinct lesion characteristics. Selective features of LM were tissue infiltration of myeloid cells producing pro-lymphangiogenic factors during early stages of active vascular growth, which occurred concomitant with an increase in cytokine levels and expansion of an immunoregulatory capillary LEC subtype, iLEC, identified through single cell transcriptomics. Importantly, macrophage depletion using colony stimulating factor 1 receptor (CSF1R) blockade, or anti-inflammatory cyclooxygenase-2 (COX-2) inhibition limited *Pik3ca^H1047R^*-driven lymphangiogenesis in mice. These results show that paracrine immune activation driven by LEC-autonomous oncogenic p110α-PI3K signaling critically contributes to pathological vascular growth in LM and provides a therapeutic target.

## Results

### Vessel type-specific responses to embryonic activation of oncogenic *Pik3ca*

Endothelial expression of *Pik3ca^H1047R^* induces excessive lymphatic vessel sprouting and localized blood vessel dilations, without sprouts, in the mouse skin (16). To explore these apparently different cellular responses of dermal lymphatic and blood ECs (LECs and BECs, respectively) to activation of PI3K signaling, we used a mouse model that allows Cre-inducible expression of *Pik3ca^H1047R^* from the ubiquitously expressed *Rosa26* locus in combination with EC-specific Cre lines (**Figure 1A**). LEC-specific *Vegfr3*-*CreER^T2^* (20) and pan-endothelial *Cdh5-CreER^T2^* (21) lines were complemented with a new transgenic mouse model that allows BEC-specific expression of *CreER^T2^* under the control of *Flt1* (encoding VEGFR1) promoter (**Supplemental Figure 1A**). Flow cytometry and immunostaining analyses confirmed that the *Vegfr1-CreER^T2^* transgene drives efficient recombination of the *R26-mTmG* reporter allele and green fluorescent protein (GFP) expression upon tamoxifen administration specifically in BECs in the skin (**Supplemental Figure 1, B-E**).

**Figure 1.**
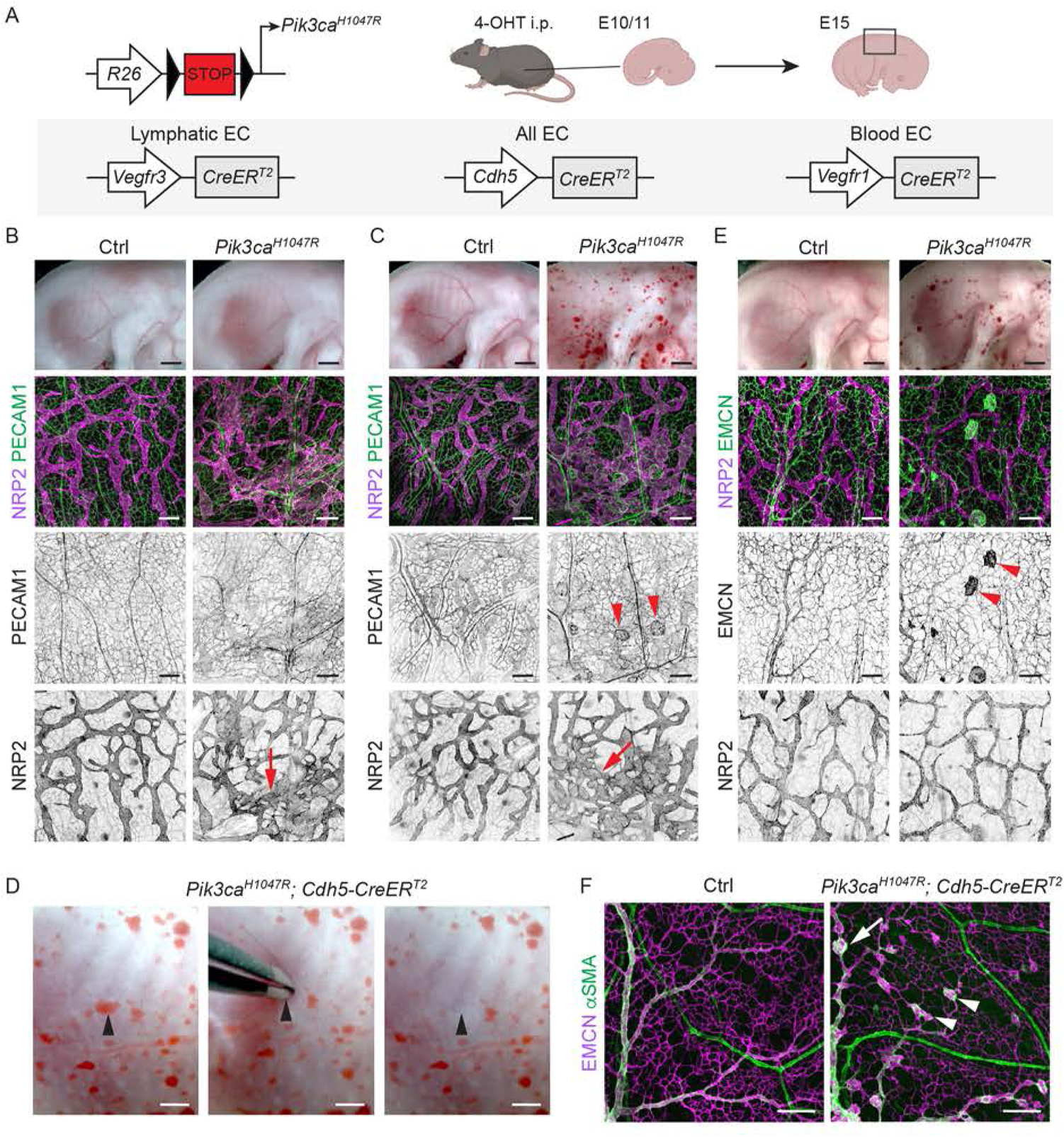
Vessel-type specific responses to activation of oncogenic *Pik3ca* signaling in the embryonic vasculature. (**A**) Genetic constructs and strategy for tamoxifen-inducible *Pik3ca^H1047R^* expression in embryonic lymphatic (*Vegfr3-CreER^T2^*), lymphatic and blood (*Cdh5-CreER^T2^*), or specifically in blood (*Vegfr1-CreER^T2^*) endothelia. (**B**, **C**) E15 *Pik3ca^H1047R^*;*Vegfr3-CreER^T2^* (**B**), *Pik3ca^H1047R^*;*Cdh5-CreER^T2^* (**C**), and their littermate control (Ctrl) embryos treated with 4-OHT at E11. Whole-mount immunofluorescence of the back skin is shown below. Single channel images show hyperbranching of NRP2^+^ lymphatic vessels (red arrows) in both models, but presence of PECAM1^+^ blood vessel lesions (red arrowheads) only in the *Cdh5-CreER^T2^* model. (**D**) Evacuation of blood upon application of pressure on a blood-filled lesion (arrowhead) in the skin of E15 *Pik3ca^H1047R^*;*Cdh5-CreER^T2^* embryo. (**E**) E15 *Pik3ca^H1047R^*;*Vegfr1-CreER^T2^* embryos treated with 4-OHT at E10, and whole-mount immunofluorescence of the back skin showing EMCN^+^ lesions (red arrowheads) in the blood vessels and normal lymphatic vasculature. (**F**) Whole-mount immunofluorescence of E17 skin from *Pik3ca^H1047R^*;*Cdh5-CreER^T2^* and littermate control (Ctrl) embryos treated with 4-OHT at E14. Note that lesions are present in the EMCN^+^ veins (arrow) and capillaries (arrowheads) but not in the αSMA^+^ arteries. Scale bar: 1 mm (B, C, E, top panels) 200 µm (B-F).

To mimic the congenital *PIK3CA*-driven vascular malformations, we first induced *Pik3ca^H1047R^* expression in LECs and/or BECs during embryonic development by administering 4-hydroxytamoxifen (4-OHT) to pregnant females at embryonic day (E) 10 or 11 (**Figure 1A**). As previously reported (16), *Vegfr3*-*CreER^T2^*-driven activation of *Pik3ca^H1047R^* expression led to hypersprouting of neuropilin-2 (NRP2)^+^ lymphatic vessels in the thoracic skin of E15 embryos while blood vessels were not affected (**Figure 1B**). Pan-EC specific expression of *Pik3ca^H1047R^* similarly resulted in a hyperbranched lymphatic vasculature, but also in multiple blood-filled lesions (**Figure 1C**) that were connected to the blood circulation as evidenced by evacuation of blood upon application of pressure (**Figure 1D**). BEC-specific activation of *Pik3ca^H1047R^* expression using the *Vegfr1*-*CreER^T2^* line at E10 led to formation of blood vessel lesions that resembled those in the *Cdh5*-*CreER^T2^* embryos, but did not affect the lymphatic vasculature (**Figure 1E**). Whole-mount immunofluorescence staining of embryonic back skin showed localized vessel dilations that were positive for the pan-endothelial marker PECAM1 (**Figure 1C**) and the venous/capillary EC marker Endomucin (EMCN) (**Figure 1E**). In contrast, EMCN negative and alpha smooth muscle actin (αSMA) positive arteries were not affected (**Figure 1F**).

Taken together, these results demonstrate that chronic activation of p110α signaling triggers a distinct response in different dermal vessel types. Lymphatic capillaries in mutant embryos expand by sprouting, whereas blood capillaries and veins show localized vessel dilations, but arteries are not affected.

### Distinct EC-autonomous responses to oncogenic *Pik3ca* underlie vessel-type specific lesion morphology

To allow analysis of the step-by-step development of *Pik3ca^H1047R^*-driven lymphatic and vascular overgrowth, we utilized postnatal mouse ear skin as a model (16). Cre-mediated recombination was induced in 3-week-old *Vegfr3-CreER^T2^* and *Vegfr1-CreER^T2^* mice by topical application of 4-OHT (**Figure 2A**). To first assess the specificity of Cre-mediated recombination, we analyzed transgenic mice carrying the *R26-mTmG* reporter allele. Efficient induction of GFP expression was observed in the ear skin vasculature with lower frequency of GFP^+^ ECs in other analyzed tissues (**Figure 2B, Supplemental Figure 2, A and B**), indicating locally restricted recombination as opposed to tissue-wide recombination observed upon systemic 4-OHT administration ((20) and data not shown). As expected, recombination was EC-subtype specific such that dermal LECs were specifically targeted in the *Vegfr3-CreER^T2^* mice (**Figure 2B, Supplemental Figure 2A**) and BECs in the *Vegfr1-CreER^T2^* mice (**Figure 2B, Supplemental Figure 2B**).

**Figure 2.**
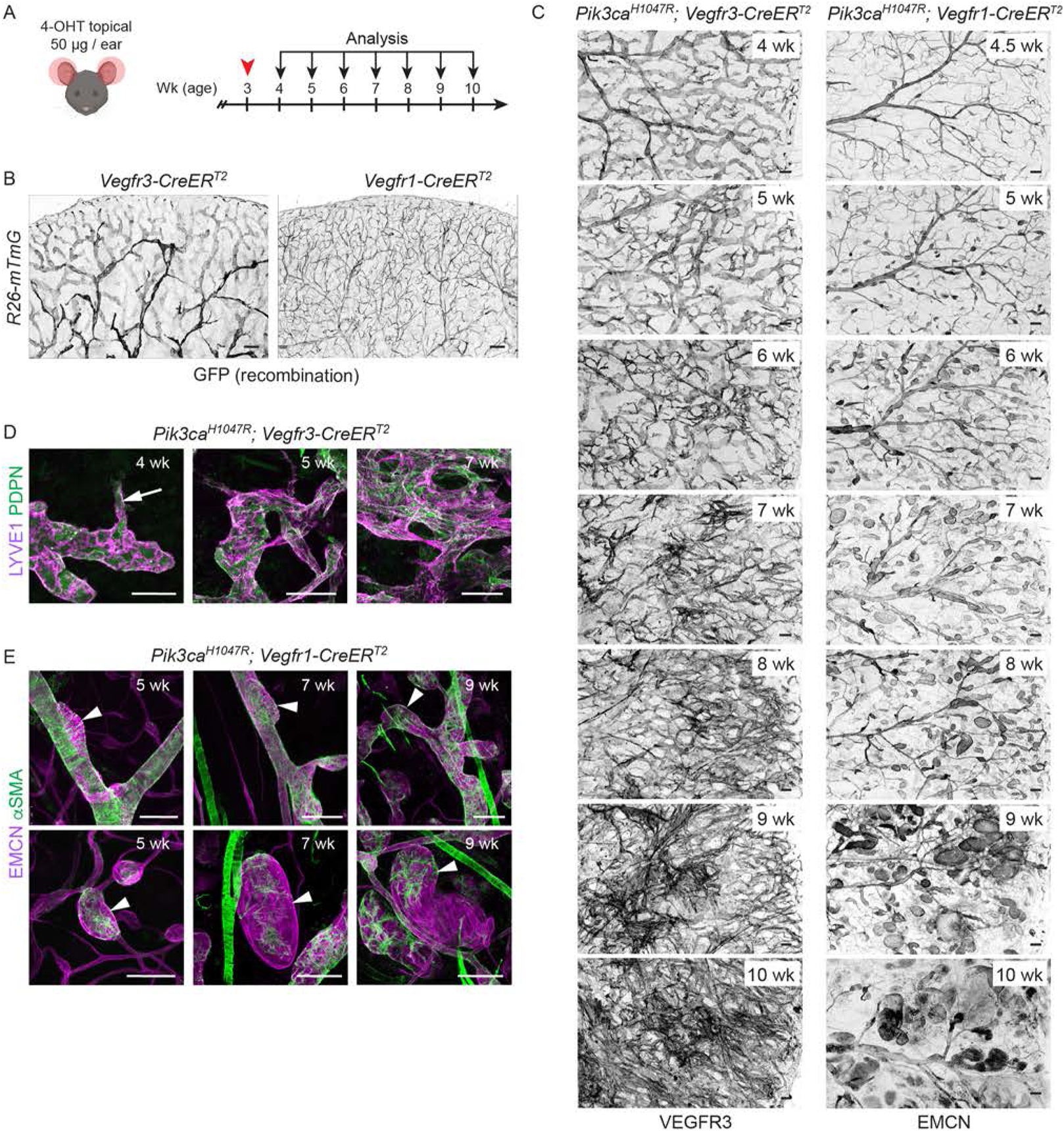
Distinct endothelial cell-autonomous responses to oncogenic *Pik3ca* in lymphatic and blood vessels. (**A**) Experimental scheme for postnatal induction of *Pik3ca^H1047R^*-driven vascular overgrowth in the dermal vasculature. (**B**, **C**) Whole-mount staining of ears from 4-OHT-treated *Vegfr3-CreER^T2^* and *Vegfr1-CreER^T2^* mice in combination with the *R26-mTmG* reporter (**B**), or the *Pik3ca^H1047R^* transgene (**C**), analyzed at the indicated stages after induction. Note efficient and EC-type specific recombination (GFP expression), and progressive *Pik3ca^H1047R^*-driven vascular overgrowth in both models. (**D**, **E**) Whole-mount immunofluorescence of the ear skin showing the formation of vessel sprouts (arrow) and hyperbranched lymphatic vessel network in the *Pik3ca^H1047R^*;*Vegfr3-CreER^T2^* mice (**D**), as opposed to vessel dilations without sprouts (arrowheads) in veins (upper panels) and venules (lower panels) of *Pik3ca^H1047R^*;*Vegfr1-CreER^T2^* mice (**E**). Note ectopic coverage by αSMA^+^ SMCs of the small lesions. Scale bar: 200 µm (B, C), 100 µm (D, E).

Next, we analyzed the progression of the vascular phenotype upon LEC- or BEC-specific induction of *Pik3ca^H1047R^* expression up to seven weeks after 4-OHT administration (i.e., at 10 weeks of age) (**Figure 2A**). In agreement with previous data (16), *Vegfr3-CreER^T2^*-driven expression of *Pik3ca^H1047R^* induced the formation of lymphatic sprouts, which progressively developed into a dense hyperbranched vessel network (**Figure 2, C and D, Supplemental Figure 3, A and C**). In contrast, and similar to the embryonic skin, ear skins of *Pik3ca^H1047R^*;*Vegfr1-CreER^T2^* mice showed localized vessel dilations in both EMCN^+^ veins and smaller EMCN^+^ venules and capillaries (**Figure 2, C and E, Supplemental Figure 3, B and C**). The lesions progressively increased in number and size, in particular in the smaller caliber vessels (**Figure 2, C and E, Supplemental Figure 3, B and C**). EMCN^−^ αSMA^+^ arteries were not affected, but we observed abnormal coverage of the capillary/venous-derived lesions by αSMA^+^ smooth muscle cells (SMCs) (**Figure 2E**). Disorganized SMC coverage is a hallmark of human VM and thought to result from impaired SMC recruitment (9). Analysis of the initial stages of lesion formation in the postnatal (**Figure 2E**) and embryonic (**Supplemental Figure 4, A and B**) skin revealed that lesions originating from larger veins with pre-existing coverage by SMCs induced disruption of the muscle layer at the site of vascular overgrowth. In the normal E17 skin αSMA^+^ SMCs are, however, absent within the capillary bed (**Supplemental Figure 4A**). Lesions within the capillary bed thus formed at a distance from αSMA^+^ vessels and instead showed ectopic recruitment and progressive increase in the number of αSMA^+^ SMCs, concomitant with increased deposition of extracellular basement membrane (**Supplemental Figure 4, A and B**).

In conclusion, the analyses of the early stages of lesion formation demonstrate different EC-autonomous responses induced by activation of p110α signaling that underlie the vessel-type specific lesion morphologies. They further suggest that certain lesion characteristics associated with *Pik3ca*-driven VM, such as the disorganized SMC layer, may develop through different mechanism depending on the origin of the lesion within the vascular network.

### Oncogenic *Pik3ca* promotes sustained EC proliferation in the lymphatic vasculature

Vascular malformations are characterized as benign, non-proliferative vessel anomalies. However, since *Pik3ca*-driven LM and VM are somatic diseases, the initial stage of lesion formation likely involves proliferation and selective expansion of the mutant ECs. In support of this, flow cytometry analysis showed an increase in the frequency of ECs expressing the proliferation marker protein Ki67 in the *Pik3ca^H1047R^;Cdh5-CreER^T2^* ear skin, which was apparent already one week after 4-OHT administration and increased after two weeks (**Supplemental Figure 5A**). Quantitative RT-PCR analysis of ECs sorted by FACS from the ears of 5-week-old *Pik3ca^H1047R^;Cdh5-CreER^T2^* mice confirmed upregulation of *Mki67* (encoding Ki67) in mutant LECs and BECs compared to controls (**Supplemental Figure 5B**).

Next, we performed whole-mount immunofluorescence of the ear skin to localize the proliferating ECs within the abnormal vascular structures, and to correlate proliferation with changes in vessel morphology. *Pik3ca*-driven vascular overgrowth was induced specifically in LECs or BECs, using the previously validated mouse models (**Figure 2C**). S-phase cells were labelled by intraperitoneal injection of EdU 16 hours prior to analysis, and combined with Ki67 staining of all cycling cells. The abnormal lymphatic sprouts in the *Pik3ca^H1047R^;Vegfr3-CreER^T2^* mice frequently contained proliferating LECs (**Figure 3A**). Quantification of the frequency of PROX1^+^LYVE1^+^ LECs that were positive for EdU and/or Ki67 revealed a high proliferation rate one week after 4-OHT administration that was sustained at ~2-fold higher level compared to control up to at least seven weeks post-induction (i.e. 10 weeks of age) (**Figure 3B**). A similar proliferative response was observed in the BECs within the developing lesions of EMCN^+^ veins and venules of *Pik3ca^H1047R^;Vegfr1-CreER^T2^* mice during the first two weeks after 4-OHT induction (**Figure 3, C and D**). However, six weeks after 4-OHT induction, i.e. at nine weeks of age, BEC proliferation rate in the lesions was reduced to that of controls (**Figure 3, C and D**). Flow cytometry analysis of *Pik3ca^H1047R^;Vegfr3-CreER^T2^* ear skin confirmed a sustained increase in the frequency of Ki67^+^ LECs (**Figure 3E**), and consequently a dramatic increase in the proportion of LECs of the total dermal EC population (**Figure 3F**). In contrast, Ki67^+^ BECs were observed only at the early stage of vascular lesion formation in the *Pik3ca^H1047R^;Vegfr1-CreER^T2^* mice (**Figure 3E**), resulting in a small increase in total BEC numbers (**Figure 3F**).

**Figure 3.**
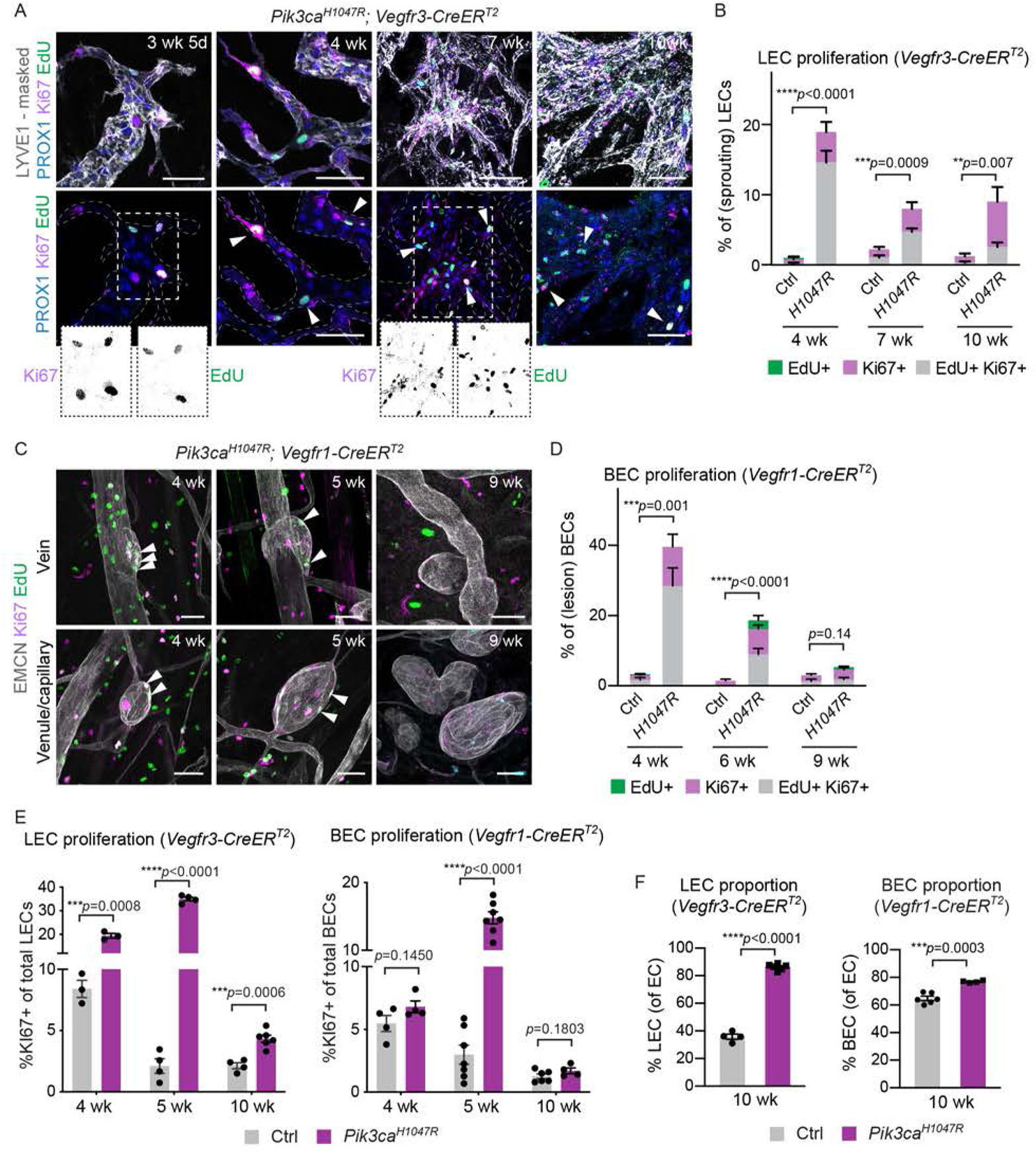
Different proliferation dynamics of LECs and BECs during *Pik3ca*-driven vascular overgrowth. (**A-D**) Whole-mount immunofluorescence of ear skin analyzed at different stages after 4-OHT administration, and quantification of S-phase cells (EdU^+^) and all cycling cells (Ki67^+^) (arrowheads) in *Pik3ca^H1047R^;Vegfr3-CreER^T2^* (**A**, **B**) and *Pik3ca^H1047R^;Vegfr1-CreER^T2^* (**C**, **D**) mice. Edu was administered 16 hours prior to analysis. LYVE1 and PROX1 were used for the identification of LECs, and EMCN for the identification of (venous) BECs. IMARIS surface mask based on LYVE1 expression was used to extract LEC-specific Ki67/EdU signals. The original unmasked images are shown in Supplemental Figure 5C. Note initial proliferative response in both models, but sustained proliferation only in the LECs of *Vegfr3-CreER^T2^* mice. (**E**) Flow cytometry analysis of proliferating dermal LECs and BECs in *Pik3ca^H1047R^;Vegfr3-CreER^T2^* and *Pik3ca^H1047R^;Vegfr1-CreER^T2^* mice, respectively, and their littermate controls at the indicated stages. Data represent mean % of Ki67^+^ ECs (*n*=3-7 mice) ± s.e.m. (**F**) Proportion of LECs and BECs out of all ECs in the *Pik3ca^H1047R^;Vegfr3-CreER^T2^* and *Pik3ca^H1047R^;Vegfr1-CreER^T2^* animals analyzed in (**E**). *p* value in (B, D, E, F), Two-tailed unpaired Student’s t-test. Scale bar: 50 μm (A, C).

In summary, the above data demonstrate that the initial stage of *Pik3ca*-driven vascular pathology involves increased EC proliferation both in lymphatic and blood vessels, likely through cell-autonomous mechanisms. However, in advanced lesions the proliferation of BECs ceased whereas LEC proliferation was sustained.

### *Pik3ca*-driven LM is associated with increased myeloid cell infiltrate

To investigate the mechanisms underlying the sustained proliferation of LECs in advanced LM lesions, we focused on the potential contribution of the immune infiltrate as a source of pro-lymphangiogenic factors such as VEGF-C (22, 23). Increased abundance of CD45^+^ cells was observed in the ear skin of *Pik3ca^H1047R^;Vegfr3-CreER^T2^* mice already one week after induction of vascular overgrowth (**Figure 4A**). In contrast, there was no apparent increase in CD45^+^ cells around the vascular lesions in *Pik3ca^H1047R^;Vegfr1-CreER^T2^* mice (**Figure 4B**). Staining for F4/80 confirmed an increased presence of macrophages, constituting the majority of dermal CD45^+^ cells (24), in the *Vegfr3-CreER^T2^*(**Figure 4, C and D**) but not in the *Vegfr1-CreER^T2^* (**Figure 4, C and E**) ears two weeks after 4-OHT induction. Flow cytometry analysis of innate and adaptive immune cells further revealed increase in the frequency (**Figure 4F**) and number (**Supplemental Figure 6A**) of CD45^+^CD11b^+^F4/80^+^ macrophages, and in particular proinflammatory Ly6C^+^ monocytes (**Figure 4F**), in *Pik3ca^H1047R^;Vegfr3-CreER^T2^* mice compared to controls. The frequency of dendritic cells (CD45^+^CD11b^+^CD11c^+^), neutrophils (CD45^+^CD11b^+^Ly6G^+^), natural killer cells (CD45^+^CD3^−^NK1.1^+^) (**Figure 4F**), T cells (CD45^+^CD3^+^CD4/CD8^+^) and B cells (CD45^+^CD3^−^B220^+^NK1.1^−^) (**Figure 4G**) were not altered at this stage. Advanced lesions, analyzed by FACS at 10 weeks of age (7 weeks after 4-OHT administration) showed a sustained increase in proinflammatory Ly6C^+^monocytes (**Figure 4F**), and additionally an increased frequency of T and B cells in advanced lesions (**Figure 4G**). Immunofluorescence staining confirmed an increased abundance of F4/80^+^ macrophages in the *Pik3ca^H1047R^;Vegfr3-CreER^T2^* ears until the analysis at 10 weeks of age (**Supplemental Figure 6B**).

**Figure 4.**
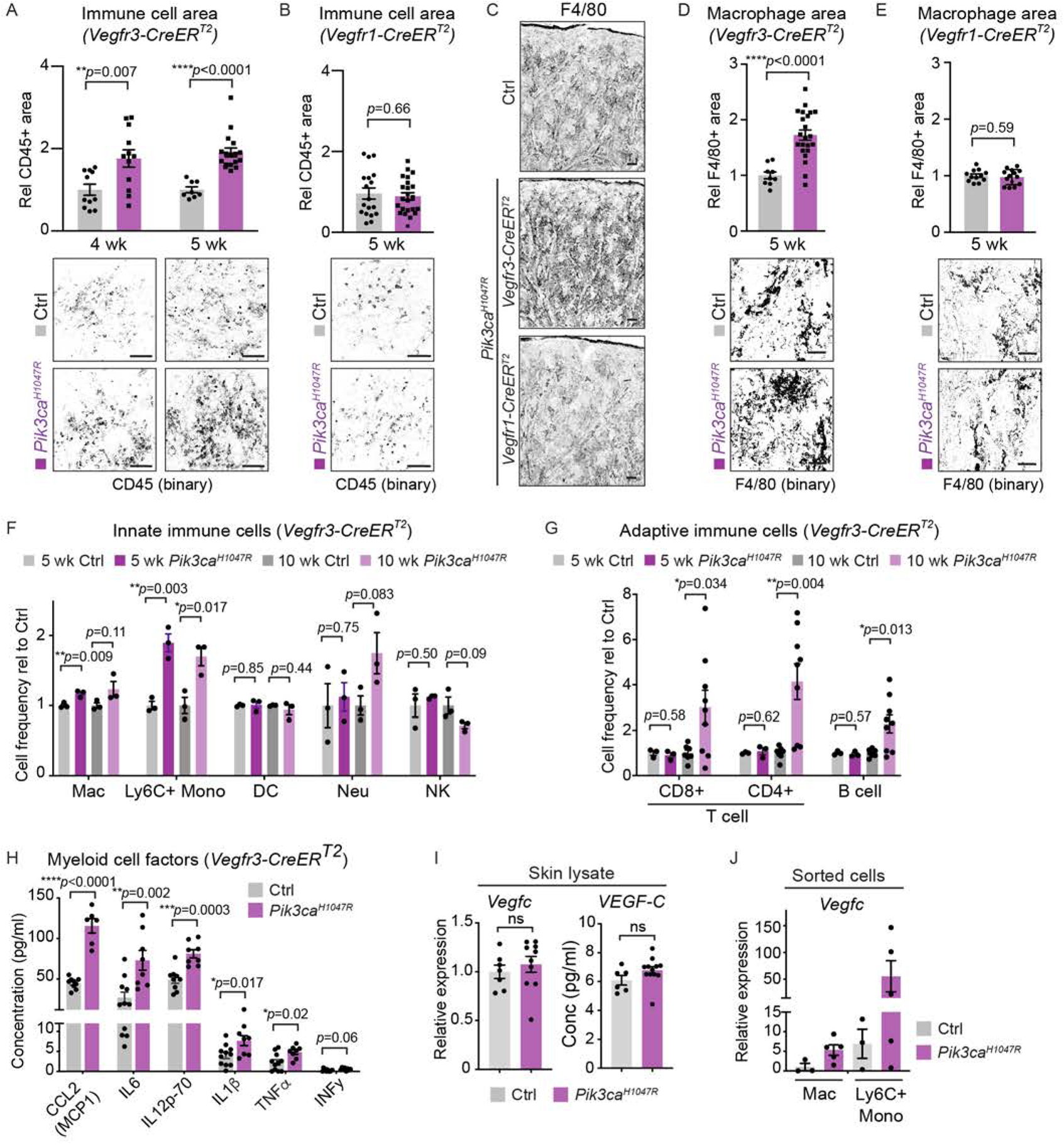
Increased inflammatory cell infiltration and pro-inflammatory cytokine levels in the *Pik3ca*-driven LM. (**A-E**) Quantification of the CD45^+^ area (**A**, **B**) and F4/80^+^ area (**C**-**E**) in the ear skin showing increase in *Pik3ca^H1047R^;Vegfr3-CreER^T2^* (**A, C, D**) but not in *Pik3ca^H1047R^;Vegfr1-CreER^T2^* (**B, C, E**) mice. Data represent mean (CD45: *n*=4-6 images from *n*=3-5 mice per genotype; F4/80: *n*=3-7 images from *n*=2-3 mice per genotype) ± s.e.m. Representative binary images are shown below the graphs. (**F, G**) Flow cytometry analysis of innate (**F**) and adaptive (**G**) immune cells in the ear skin of 4-OHT-treated 5-week-old and 10-week-old *Pik3ca^H1047R^; Vegfr3-CreER^T2^* mice and littermate controls. Mac, macrophage; Mono, monocyte; DC, dendritic cell; Neu, neutrophil; NK, natural killer cell. Data represent relative cell frequency (of live cells) relative to the control (*n*=3-9 mice) ± s.e.m. (**H**) Multiplex ELISA analysis of pro-inflammatory cytokines and chemokines associated with recruitment and/or activation of myeloid cells in whole ear skin lysates from *Pik3ca^H1047R^;Vegfr3-CreER^T2^* mice and littermate controls. Data represent mean protein levels relative to control (*n*=6-11 mice) ± s.e.m. (**I**) qRT-PCR analysis (left) and ELISA analysis (right) of *Vegfc*/VEGF-C levels in the ear skin lysates of 4-OHT-treated 5-week-old *Pik3ca^H1047R^;Vegfr3-CreER^T2^* and littermate control mice. qRT-PCR data represent mean relative expression (normalized to *Hprt*; *n*=7-10 mice) ± s.e.m. Transcript and protein levels are presented relative to controls. (**J**) qRT-PCR analysis of *Vegfc* in CD45^+^Cd11b^+^F4/80^+^ macrophages and CD45^+^Cd11b^+^F4/80^+^Ly6C^+^ monocytes, FACS-sorted from the ear skin of 4-OHT-treated 5-week-old *Pik3ca^H1047R^;Vegfr3-CreER^T2^* and littermate control mice. qRT-PCR data represent mean relative expression (normalized to *Hprt*; *n*=3-5 mice) ± s.e.m. Transcript and protein levels are presented relative to macrophages/monocytes in control mice. *p*-value in (A, B, D-I) obtained using Two-tailed unpaired Student’s t-test. Scale bar: 100 μm (A-E).

Although CD45^+^CD11b^+^F4/80^+^ myeloid cells were not significantly increased in the *Pik3ca^H1047R^;Vegfr1-CreER^T2^* mice compared to controls at 5 weeks of age (**Supplemental Figure 6C**), advanced venous lesions at 10 weeks of age showed an increased frequency of other immune populations except for T cells, and in particular myeloid cell populations, neutrophils and B cells (**Supplemental Figure 6D**). This delayed immune response may be secondary to disruption of vessel integrity and leakage in this model.

### Increased pro-inflammatory cytokine levels in *Pik3ca*-driven LM

To further assess the inflammatory status of the ears, we performed multiplex ELISA that allows simultaneous measurement of multiple pro-inflammatory cytokines and chemokines. The levels of pro-inflammatory proteins associated with the recruitment and/or activation of antigen-presenting myeloid cells, including CCL2 (also known as monocyte chemoattractant protein MCP1), IL1β, IL6 and IL12, were significantly increased in ear skin lysates of 5-week-old *Pik3ca^H1047R^;Vegfr3-CreER^T2^* mice (**Figure 4H**), but not of *Pik3ca^H1047R^;Vegfr1-CreER^T2^* mice (**Supplemental Figure 6E**). Proteins associated with the recruitment and/or activation of B cells (IL4, IL5) or T cells (IL2, IL9, IL15, IL17A) were unaltered in both models (**Supplemental Figure 6, E and F**). The major pro-inflammatory cytokines TNFα and INFγ were also unaltered in the blood sera of *Pik3ca^H1047R^;Vegfr3-CreER^T2^* mice (**Supplemental Figure 6G**), thereby excluding systemic inflammation. TNFα levels were increased in the mutant in comparison to control ear skin lysate (**Figure 4H**), but the low concentration likely reflects low-grade local chronic inflammation (25).

We additionally measured VEGF-C protein and *Vegfc* transcript levels in total ear skin lysates using ELISA and qRT-PCR analysis, respectively, and observed a trend toward higher VEGF-C levels in the mutant in comparison to the control skin (**Figure 4I**). qRT-PCR analysis of FACS-sorted CD45^+^CD11b^+^Ly6G^−^F4/80^+^ macrophages and Ly6C^+^ monocytes also showed higher *Vegfc* levels in both cell populations in mutant in comparison to control mice (**Figure 4J**). In particular, we observed a 3-20-fold increase in *Vegfc* levels in Ly6C^+^ cells in 3 out of 5 mutant ears in comparison to control ears.

Collectively, the above data demonstrate that the infiltration of VEGF-C-producing CD45^+^CD11b^+^F4/80^+^ myeloid cells and the upregulation of pro-inflammatory cytokines and chemokines aiding their recruitment are selective features of the first weeks of active lymphatic overgrowth driven by *Pik3ca^H1047R^* mutation.

### Single cell transcriptomics identifies a molecularly distinct dermal capillary LEC subtype with putative immunoregulatory functions

The pro-inflammatory molecules specifically upregulated in the *Pik3ca^H1047R^;Vegfr3-CreER^T2^* skin are expressed in a variety of cell types, including the infiltrating myeloid cell themselves (26). To determine the potential contribution of LEC-autonomous PI3K signaling in promoting a pro-inflammatory environment, we applied single-cell RNA sequencing (scRNAseq). LECs from the ear skin of *Pik3ca^H1047R^;Cdh5-CreER^T2^* (n=5) and Cre^−^ littermate (n=2) mice were subjected to scRNAseq using Smart-Seq2 (27) two weeks after 4-OHT treatment (**Figure 5A**). A separately bred untreated wild type (C57BL/6J) mouse was also included to control for possible effects of the treatment in littermate controls.

**Figure 5.**
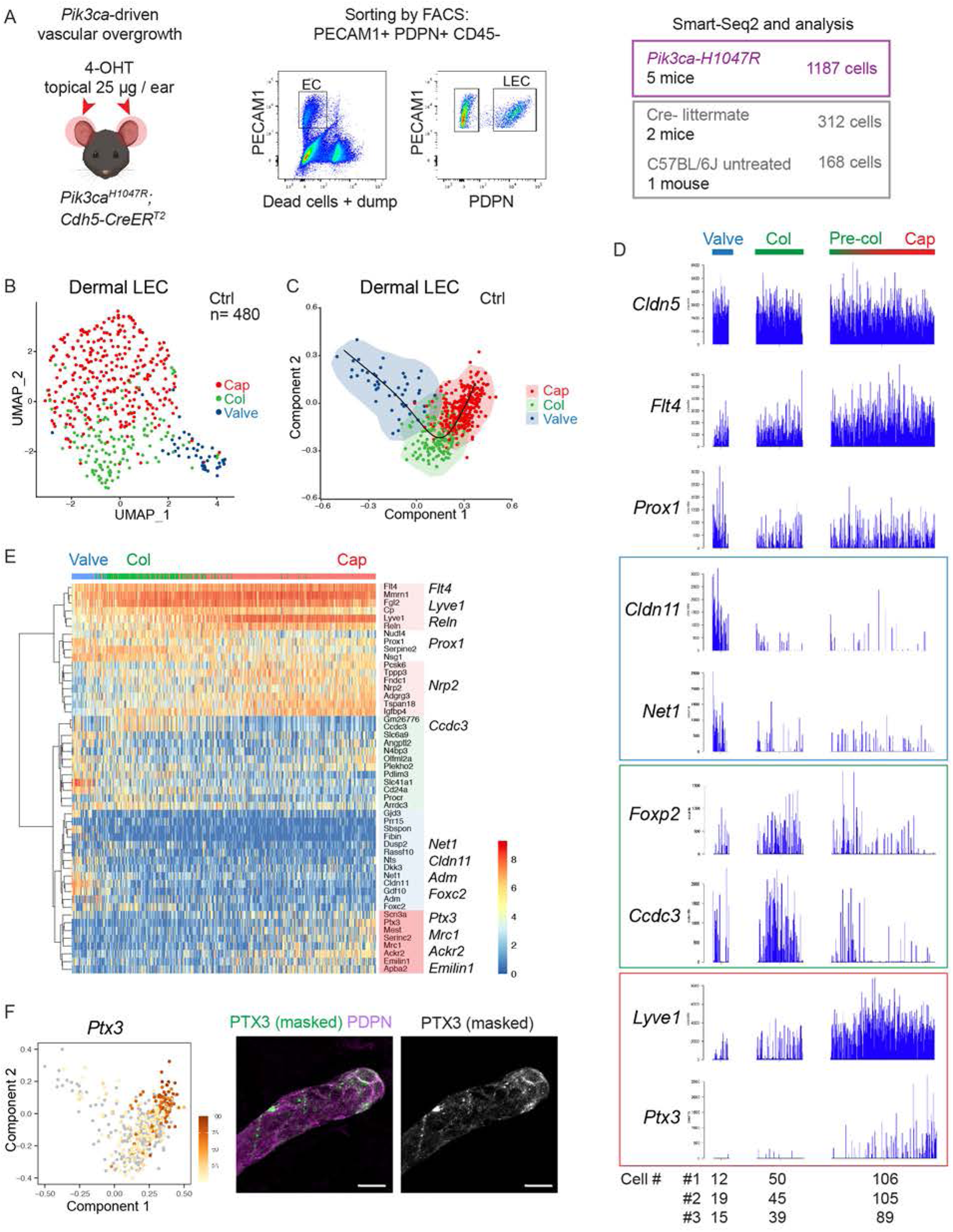
Definition of dermal LEC subtypes by single cell transcriptomics. (**A**) Schematic overview of dermal LEC isolation for single cell RNA sequencing (scRNA-seq). (**B**, **C**) The three dermal LEC clusters from control ear skin visualized in a UMAP landscape (**B**) and after trajectory inference analysis (**C**), labeled by cluster assignment. (**D)** ScRNA-seq bar plots showing the expression of selected LEC and LEC subtype marker genes in the three LEC clusters, Cells were ordered by the trajectory. Cell numbers from the different samples (mice) are shown for each cluster below the graph. (**E**) Heatmap showing the expression of zonation markers across the three LEC clusters. Cells were ordered by the trajectory. Color indicates read counts in log-scale. (**F**) *Ptx3* expression visualized in a UMAP (left) and whole-mount immunofluorescence of non-permeabilized ear skin showing extracellular PTX3 deposition predominantly at capillary terminals (right panels). IMARIS surface mask based on PDPN expression was used to extract LEC-specific PTX3 signal. The original unmasked image is shown in Supplemental Figure 7A. Scale bar: 20 μm (F).

To first define the normal transcriptome of dermal LECs, we analyzed 480 single cell transcriptomes from control mice that passed the quality controls. The cells distributed into 3 clusters after applying canonical correlation analysis (CCA) method for batch correction and Seurat graph-based clustering approach (28), and visualization using the Uniform Manifold Approximation and Projection (UMAP (29)) (**Figure 5B**). Trajectory inference analysis ordered the cells along a linear trajectory across the clusters based on similarities in their expression patterns (**Figure 5, C and D**). As expected, all clusters were characterized by high expression of pan-endothelial (*Cldn5*) and LEC-specific marker genes (*Flt4*, *Prox1*) (**Figure 5D**), and lack of BEC marker expression (*Flt1*, *Nrp1*) (data not shown). Each cluster included an equal contribution of cells from the 3 control samples (**Figure 5D**). Based on the expression of known LEC subtype markers, the clusters were annotated as valve LECs (*Cldn11^+^*) (30), collecting vessel LECs (*Foxp2^+^*) (31), and capillary LECs (*Lyve1^high^*) (**Figure 5, B-D**). Trajectory analysis based on gene expression data suggested phenotypic progression from collecting vessel to capillary LECs (**Figure 5, C-E**), which mirrors anatomical positioning of so called pre-collecting vessels that share molecular and functional features of the two vessel types (32). This analysis also identified a subset of capillary LECs (9% (42 out of 480) LECs) at the end of the trajectory that selectively expressed high levels of *Ptx3* and lacked *Foxp2* expression (**Figure 5, D-F**). LEC zonation markers highly expressed in *Ptx3^high^* dermal LECs include the extracellular matrix glycoprotein *Emilin1* (**Figure 5E**), which is a functionally important component of the elastic anchoring filaments in capillary LECs (33). *Ptx3^high^* LECs were also enriched in transcripts encoding key regulators of innate and adaptive immune responses including *Ptx3* itself as well as *Mrc1* and *Ackr2* (**Figure 5, D and E**). Whole-mount immunofluorescence of non-permeabilized ear skin revealed extracellular PTX3 around blunt-ended terminals of lymphatic capillaries (**Figure 5F**), thus confirming a distinct anatomical location of PTX3^+^ LECs. In addition, a subset of valves of pre-collecting vessels were PTX3^+^ (**Supplemental Figure 7A**). These cells may correspond to *Ptx3^+^* LECs scattered across the pre-collecting-capillary LEC cluster in the scRNAseq data (**Figure 5D**).

In summary, scRNAseq identifies dermal LEC hierarchy that recapitulates lymphatic vascular architecture, and defines a previously unknown molecularly distinct *Ptx3^high^* population within dermal capillary terminals as a putative immunoregulatory LEC subtype - termed here as iLEC.

### *Pik3ca^H1047R^* drives iLEC expansion

Next, we performed a similar analysis of LECs isolated from the *Pik3ca^H1047R^;Cdh5-CreER^T2^* mice. We obtained in total 1187 quality controlled single-cell transcriptomes that distributed into 6 clusters (**Figure 6A**). Based on the expression of the LEC subtype markers identified in the control skin dataset, we defined clusters of valve, collecting vessel and capillary LECs. We also observed a large population of *Ptx3^high^* capillary LECs that included two clusters representing non-proliferating and proliferating LECs, the latter recognized by their high expression of cell cycle genes (e.g. *Ccnb1*) (**Figure 6, A and B, Supplemental Figure 8A**). One cluster was characterized by an intermediate identity with expression of both capillary (e.g. *Lyve1*) and collecting vessel (e.g. *Mmrn2*) genes (**Figure 6, A and B, Supplemental Figure 8A**). Cluster level analysis thus suggested active expansion of the *Ptx3^high^* iLEC population in *Pik3ca^H1047R^* mutant mice, which was also evident in the subtype composition of LEC populations in the mutant in comparison to control skin (**Figure 6C**).

**Figure 6.**
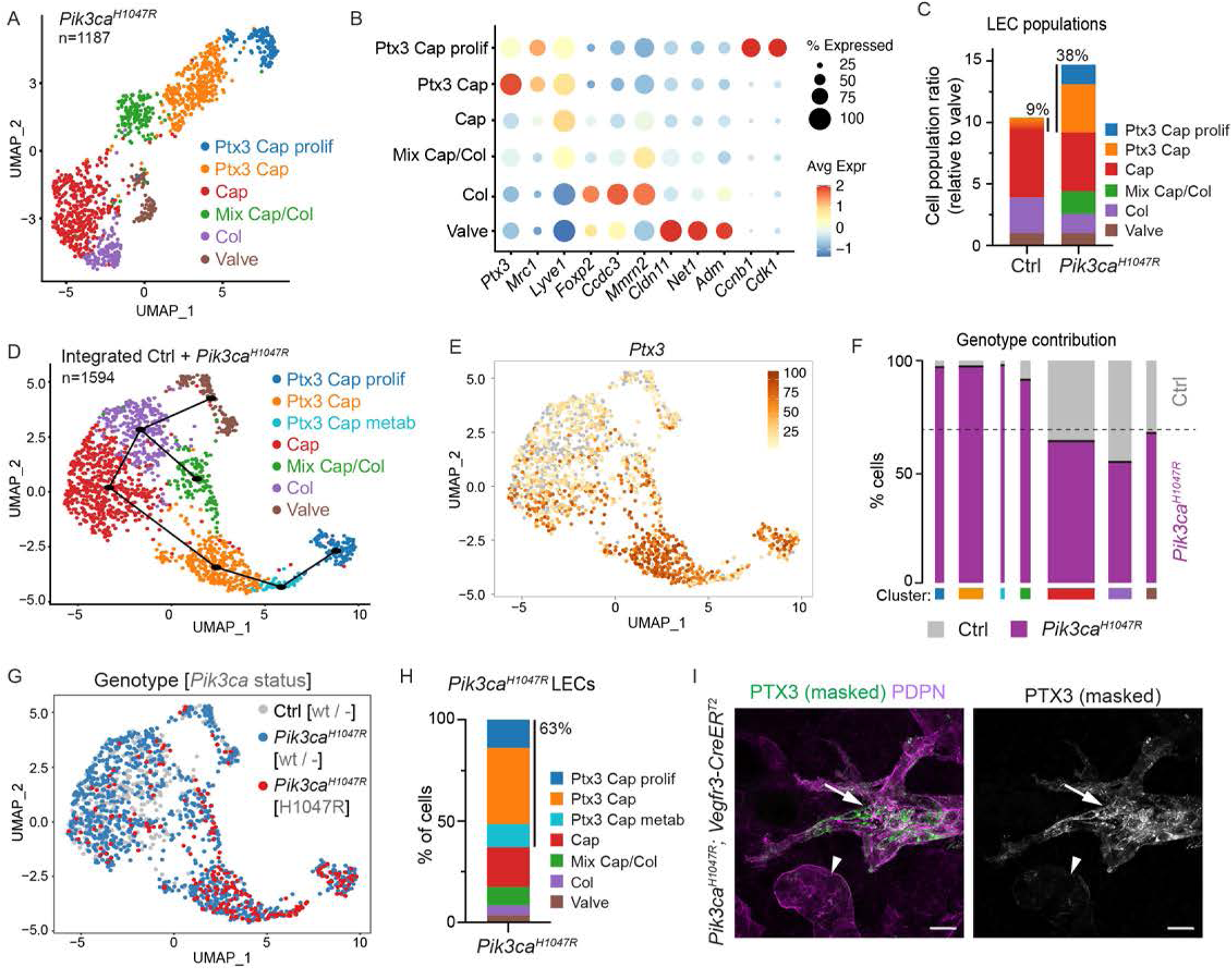
LEC-autonomous *Pik3ca* signaling induces expansion of iLECs. (**A**) The six dermal LEC clusters from *Pik3ca^H1047R^;Cdh5-CreER^T2^* ear skin visualized in a UMAP landscape. (**B**) Dot plot of LEC subtype markers for the six LEC clusters from (**A**). Dot size illustrates percentage of cells presenting transcript sequence counts and color illustrates the average expression level (log2 fold change) within a cluster. (**C)** Subtype composition of LEC populations in the control and *Pik3ca^H1047R^* mutant skin. Population sizes are shown relative to valve LECs (=1). The percentage of *Ptx3^+^* capillary LEC population of the total LEC population is indicated. (**D**) The 7 dermal LEC clusters from the dataset integrating data from control and *Pik3ca^H1047R^;Cdh5-CreER^T2^* ear skin visualized in a UMAP landscape. Black line depicts trajectories calculated from UMAP embedding of the combined dataset (unsupervised Slingshot algorithm), resulting in two branches. (**E**) *Ptx3* expression visualized in a UMAP. (**F**) Proportions of cells in the seven clusters (represented by bar width) and genotype proportions in each cluster (grey, Ctrl; purple, *Pik3ca^H1047R^*). Dashed line indicates the input levels of the two genotypes. (**G**) Genotype distribution and *Pik3ca* transcript status visualized in a UMAP, as indicated. *Ptx3^+^* capillary LEC clusters from *Pik3ca^H1047R^* mutant mice are enriched with cells expressing the mutant transcript (red). (**H**) Distribution of LECs expressing the mutant *Pik3ca^H1047R^* transcript showing the majority (63%) within the three Ptx3^high^ clusters. (**I**) Whole-mount immunofluorescence of *Pik3ca^H1047R^;Vegfr3-CreER^T2^* ear skin showing high expression of PTX3 in the abnormal lymphatic sprouts (arrows) compared to morphologically normal lymphatic capillaries (arrowheads). IMARIS surface mask based on PDPN expression was used to extract LEC-specific signals. The original unmasked images are shown in Supplemental Figure 7A. Scale bars: 20 μm (I).

We integrated the LEC single-cell transcriptomes from control and *Pik3ca^H1047R^* mutant mice (after removal of contaminants in total 1594 cells) using Harmony, and identified 7 LEC clusters (**Figure 6D**). Marker expression defined 6 clusters corresponding the same identities than those in the *Pik3ca^H1047R^* mutant mice, including non-proliferating and proliferating *Ptx3* capillary LECs (**Figure 6E, Supplemental Figure 8B**), but also an additional *Ptx3^high^* cluster characterized by high expression of metabolic genes (**Supplemental Figure 8B**). Trajectory analysis based on gene expression data suggested linear phenotypic progression between the non-proliferative and proliferative *Ptx3^high^* clusters (**Figure 6D**). Assessment of the relative contribution of cells originating from the different genotypes of mice revealed that the *Ptx3* capillary LEC clusters, as well as the cluster of mixed identity, were almost exclusively composed of LECs isolated from the *Pik3ca^H1047R^* mutant mice (**Figure 6F**). *Ptx3^high^* capillary LEC clusters further showed enrichment of cells expressing the transgene-encoded *Pik3ca^H1047R^* transcript (**Figure 6, G and H**), that was expressed at a similar level compared to the endogenous mouse *Pik3ca* transcript (**Supplemental Figure 8C**). Whole-mount immunofluorescence confirmed increased expression of PTX3 (**Figure 6I, Supplemental Figure 7A**) in the abnormal lymphatic vessel sprouts in *Pik3ca^H1047R^*;*Vegfr3-CreER^T2^* mice, further supporting the selective expansion of the *Ptx3* capillary iLEC population in the mutant skin.

### Increased PTX3 expression in human LM

To explore potential clinical relevance of the findings, we analyzed PTX3 expression in normal human skin and in lymphatic malformations. Clinical features of three LM patients with a *PIK3CA^H1047R^* mutation selected for the study are summarized in **Supplemental Table 1**. Immunofluorescence staining of paraffin sections of normal skin showed deposition of PTX3 around PDPN^+^ lymphatic vessels in control tissue but low expression in LECs (**Figure 7, A-C**). In contrast, LECs within LM lesions showed strong immunoreactivity (**Figure 7, A-C**). PTX3 immunostaining intensity, measured as Corrected Total Cell Fluorescence, was 5-fold higher in LECs from LM in comparison to control tissue (**Figure 7D**), and covered a 2-fold larger area of PDPN^+^ lymphatic vessels (**Figure 7E**).

**Figure 7.**
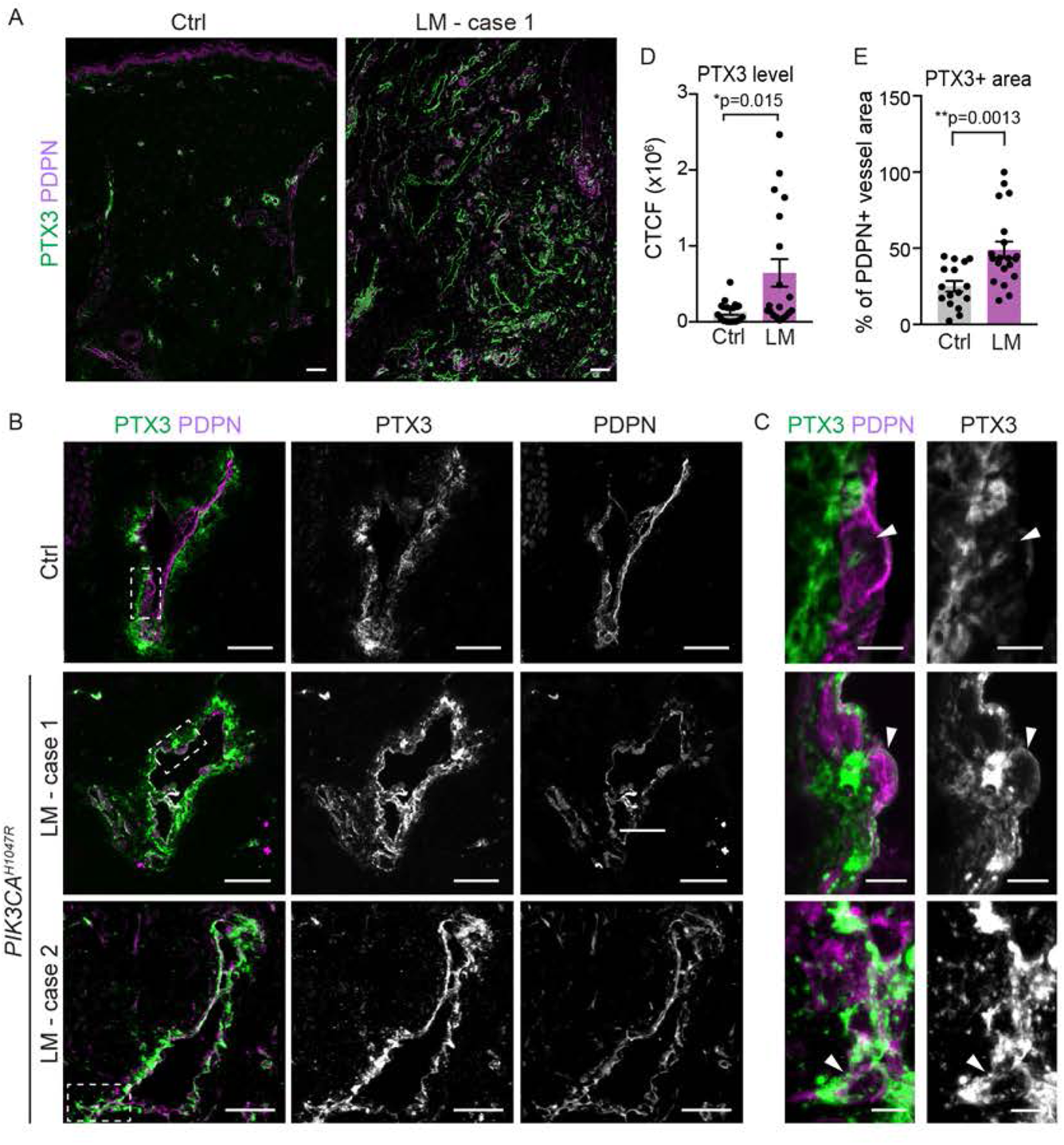
Increased PTX3 deposition and expression in human *PIK3CA^H1047R^*-driven LM. (**A-C**) Immunofluorescence staining of PTX3 in paraffin sections of normal skin and lymphatic malformations (LM) with a *PIK3CA^H1047R^* mutation. Boxed areas in (B) are magnified in (C). Note deposition of PTX3 around PDPN^+^ lymphatic vessels in control tissue but low expression in LECs, while LECs from LM tissue show PTX3 immunoreactivity (arrowheads in (C)). (**D, E**) Quantification of PTX3 immunoreactivity in PDPN^+^ lymphatic vessels showing higher intensity, measured as Corrected Total Cell Fluorescence (CTCF) (**D**), and larger PTX3^+^ vessel area (**E**) in LECs from LM in comparison to control tissue. Scale bars: 100 μm (A), 50 μm (B), 5 μm (C).

Taken together, the expansion and active proliferation of the *Ptx3^high^* capillary iLECs in the mouse model of *Pik3ca^H1047R^*-driven LM, and high lymphatic endothelial expression and deposition of PTX3 in human LM suggest PTX3^high^ LECs as pathogenic cells in these vascular lesions.

### *Pik3ca^H1047R^* induces a pro-inflammatory transcriptome in iLECs

To identify LEC-autonomous *Pik3ca*-driven transcriptional changes we next focused on the pathological *Ptx3* capillary LEC clusters representing iLECs. To avoid the confounding effect of the cell-cycle (34), we determined differentially expressed genes between the non-proliferative *Ptx3* clusters in mutant mice in comparison to *Ptx3* capillary LECs from control mice. Gene Ontology (GO) analysis of differentially expressed genes revealed enrichment of biological processes related to metabolic processes, cell cycle transition, cell migration and cell-matrix adhesion in the mutant clusters (**Supplemental Figure 9, A and B**), consistent with the established role of the PI3K pathway (10) and previously reported migratory phenotype of *Pik3ca^H1047R^* expressing LECs *in vitro* and *in vivo* (16). Both mutant clusters also showed enrichment of processes and genes related to immune regulation (**Figure 8, A and B, Supplemental Figure 9C**). The latter include upregulation of genes encoding pro-inflammatory cytokines (*Ccl2, Ccl7*), (scavenger) receptors (*Ackr2, L1cam*), as well as extracellular matrix proteins (*Lgals3*) and proteinases (*Adam17*, *Adam8*, *Mmp14, Mmp2*) implicated in inflammatory processes (**Figure 8C**). Conversely, downregulated biological processes include negative regulation of inflammatory processes (**Supplemental Figure 9C**). Lymphatic endothelial expression of CCL2/MCP1 in *Pik3ca^H1047R^*;*Vegfr3-CreER^T2^* mice was confirmed by whole-mount immunofluorescence, while no staining was detected in the control skin (**Figure 8D, Supplemental Figure 7B**). Increased CCL2/MCP1 levels observed in the mutant ear skin lysates (**Figure 4J**) are thus, at least in part, due to LEC-autonomous expression. Taken together, single cell transcriptomics revealed that *Pik3ca^H1047R^* promotes a pro-inflammatory transcriptome in LECs.

**Figure 8.**
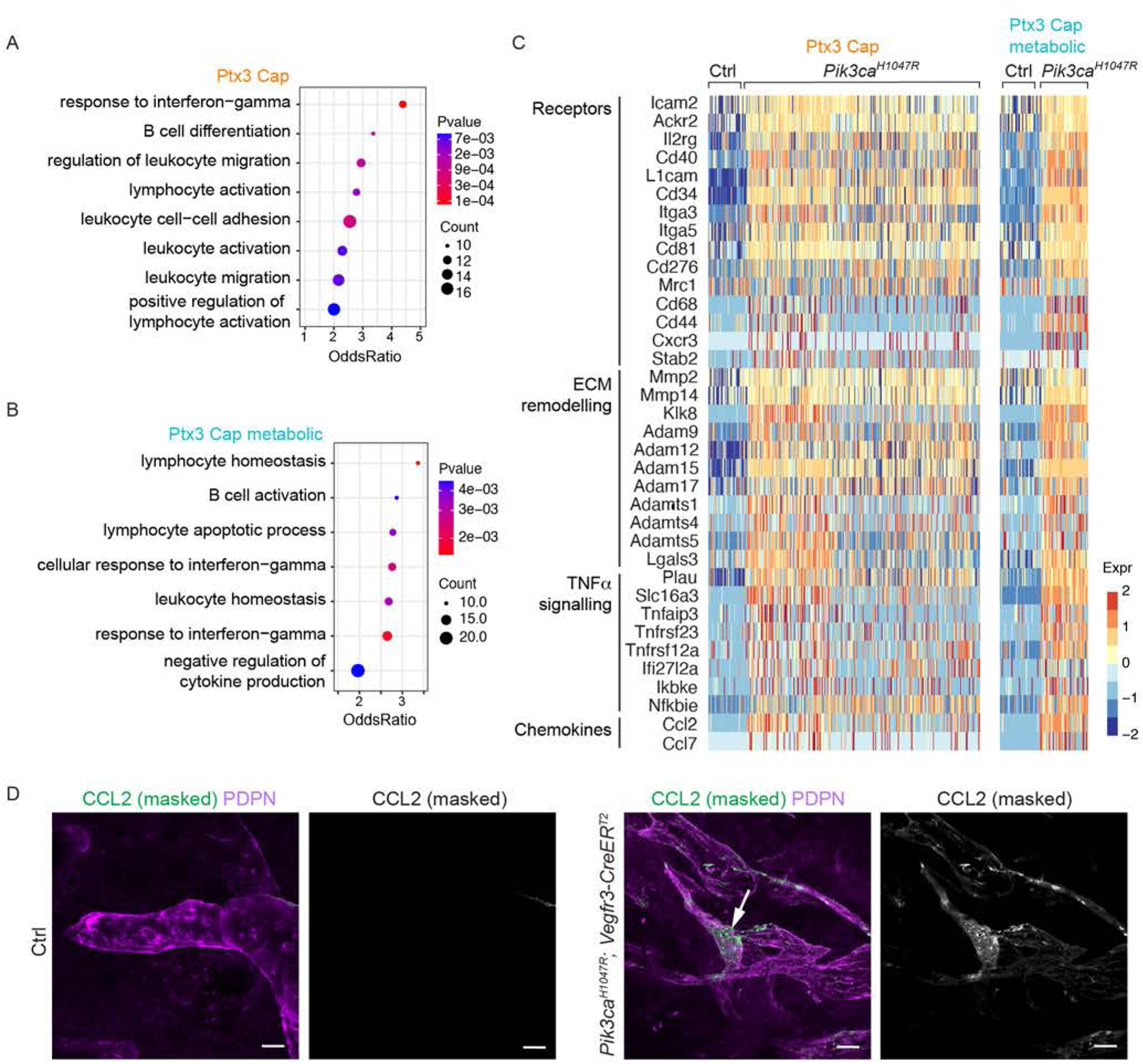
Pro-inflammatory transcriptome of *Pik3ca^H1047R^*-expressing LECs. (**A**) Gene ontology (GO) analysis of genes enriched in the clusters of non-proliferative *Ptx3^+^* capillary LECs from *Pik3ca^H1047R^* in comparison to *Ptx3^+^* capillary LECs in control mice. Selected terms for enriched immune-related biological process are shown. Dot size illustrates count number and color illustrates adjusted p-value. (**B**) Heatmap showing the expression of upregulated immune-related genes in *Ptx3^+^* capillary LECs from *Pik3ca^H1047R^* mutant in comparison to control mice. Color illustrates the expression level (log2 fold change). (**C**) Whole-mount immunofluorescence of *Pik3ca^H1047R^;Vegfr3-CreER^T2^* ear skin showing lymphatic endothelial expression of CCL2/MCP1 in the mutant. Scale bars: 20 μm (C).

### Anti-inflammatory therapy limits LM growth

Based on the increased lymphatic endothelial expression of immune regulatory molecules, we hypothesized that paracrine immune activation may critically contribute to promoting pathological lymphangiogenesis in the *Pik3ca^H1047R^* mice. To inhibit the expansion and differentiation of macrophages (35), we administered a blocking antibody against the macrophage colony-stimulating factor 1 receptor (CSF1R) from the time of induction of *Pik3ca^H1047R^* expression (**Figure 9A**). A 2.5-week treatment period with anti-CSF1R antibody partially depleted F4/80+ macrophages (**Figure 9B**) and significantly inhibited lymphatic vessel growth in the ear skin of *Pik3ca^H1047R^;Vegfr3-CreER^T2^* mice (**Figure 9, C and D**). This indicates a critical requirement of monocyte/macrophage populations in the initial stages of LM growth.

**Figure 9.**
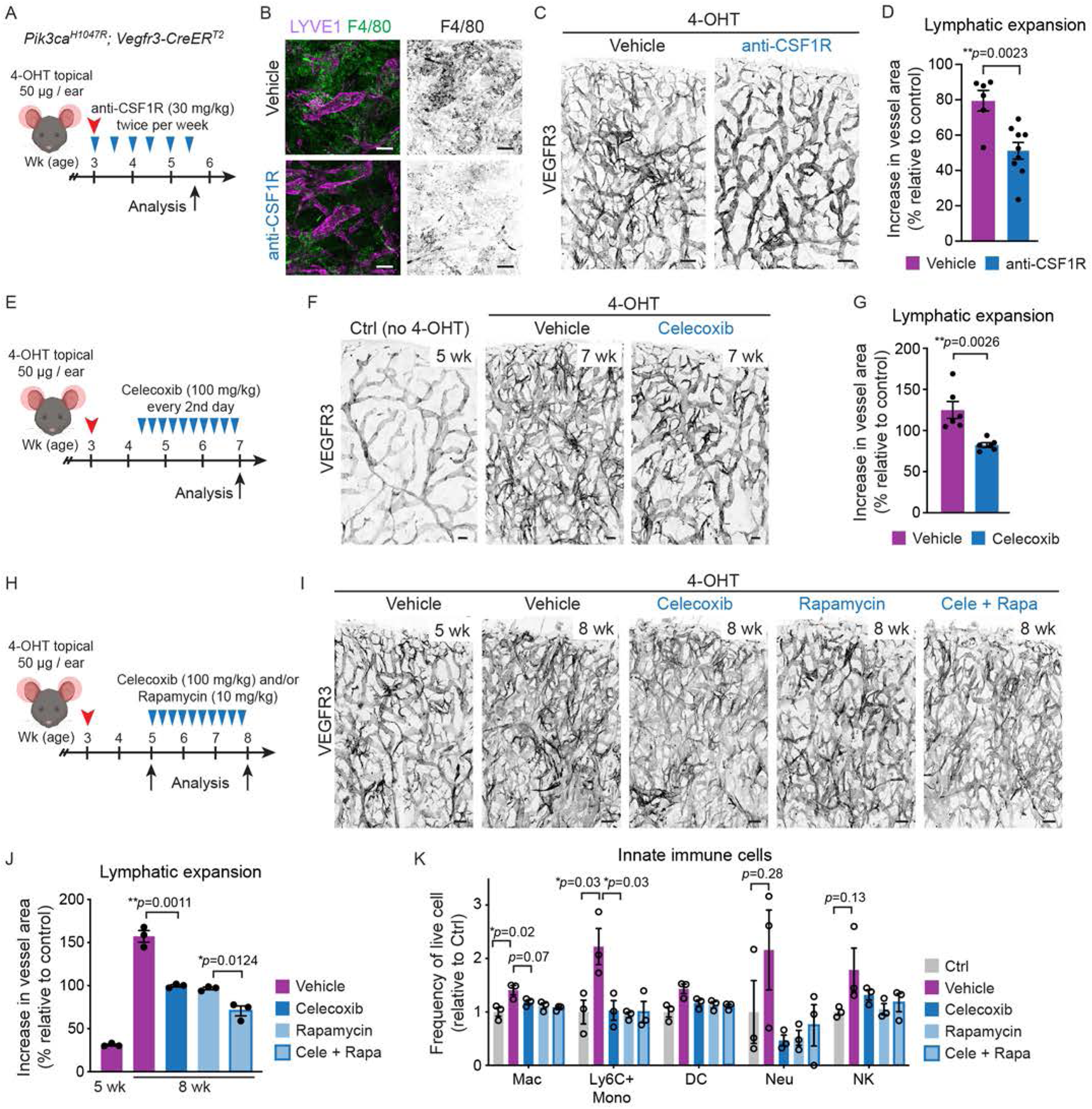
Anti-inflammatory treatment limits *Pik3ca^H1047R^*-driven lymphangiogenesis. (**A**) Experimental scheme for depletion of monocyte/macrophage populations using anti-CSF1R antibody in the mouse model of *Pik3ca^H1047R^*-driven LM. (**B, C**) Whole-mount immunofluorescence of ears from a control (Ctrl) mouse, and 5.5-week-old 4-OHT-treated *Pik3ca^H1047R^*;*Vegfr3-CreER^T2^* mice following a 2.5-week treatment with anti-CSF1R or vehicle for macrophage (B) and lymphatic markers (B, C). (**D**) Quantification of lymphatic vessel area, shown as % increase in vessel area in comparison to untreated (no 4-OHT) control. Data represent mean (*n*=6 (vehicle) or 9 (anti-CSF1R) mice) ± s.e.m. (**E**) Experimental scheme for celecoxib and/or rapamycin treatment of *Pik3ca^H1047R^*-driven LM. (**F**) Whole-mount immunofluorescence of ears from a control (Ctrl) mouse, and 7-week-old 4-OHT-treated *Pik3ca^H1047R^*;*Vegfr3-CreER^T2^* mice following a 2-week treatment with celecoxib or vehicle. (**G**) Quantification of lymphatic vessel area, shown as % increase in vessel area in comparison to control. Data represent mean (*n*=6 mice) ± s.e.m. (**H**) Experimental scheme for extended celecoxib treatment of established *Pik3ca^H1047R^*-driven LM. (**I**) Whole-mount immunofluorescence of ears from 4-OHT-treated *Pik3ca^H1047R^*;*Vegfr3-CreER^T2^* mice at the start of the treatment period (5 wk), and following a 3-week treatment with celecoxib and/or rapamycin, or vehicle (8 wk). (**J**) Quantification of lymphatic vessel area, shown as % increase in vessel area in comparison to the control. Data represent mean (*n*=3 mice) ± s.e.m. (**K**) Flow cytometry analysis of innate immune cells in the ear skin of 4-OHT-treated 8-week-old *Pik3ca^H1047R^; Vegfr3-CreER^T2^* mice and littermate controls following treatment as in (**H**). Mac, macrophage; Mono, monocyte; DC, dendritic cell; Neu, neutrophil; NK, natural killer cell. Data represent relative cell frequency (of live cells) relative to the control (*n*=3 mice) ± s.e.m. *p* value in (C, F, I), Two-tailed unpaired Student’s t-test. Scale bars: 100 μm (B, C, F, I).

Monocyte-derived macrophages are associated with the formation of lymphoid aggregates and tertiary lymphoid organs (TLOs) in chronic inflammation (36), and frequently found in patients with LMs (37), suggesting that efficient treatment of advanced lesions requires targeting of more complex inflammatory environment. Inhibition of cyclooxygenase-mediated production of prostaglandins, and in particular cyclooxygenase 2 (COX-2) using celecoxib, has a potent effect on both macrophage recruitment and cytokine release (38, 39), but also immunosuppressive effect on T cells (40) that we found to be increased in advanced LM lesions in mice. To test the therapeutic effect of COX-2 selective inhibition on LM progression, we administered celecoxib to *Pik3ca^H1047R^;Vegfr3-CreER^T2^* mice 10 days after induction of lymphatic vessel overgrowth (**Figure 9E**). Systemic COX-2 inhibition significantly reduced vessel growth after a 2-week treatment period (**Figure 9F**). Notably, the 34% reduction in *Pik3ca*-driven vascular expansion upon celecoxib treatment (**Figure 9G**) is comparable to the effect of the mTOR inhibitor rapamycin reported after a 1.5-week treatment period in this model (16).

To assess the effects of celecoxib and rapamycin on more advanced lesions and their immune cell infiltration, we administered them alone or in combination 2 weeks after induction of lymphatic vessel overgrowth and analyzed the mice after a 3-week treatment period (**Figure 9H**). Individually administered inhibitors showed a similar 60% reduction in vascular expansion, and an additive effect with 87% reduction when used in combination (**Figure 9, I and J**). FACS analysis of the ear skin of celecoxib-treated *Pik3ca* mutant mice showed a decrease in the frequency of CD45^+^ cells (**Supplemental Figure 10A**), as well as F4/80^+^ macrophages and Ly6C^+^ proinflammatory monocytes (**Figure 9K**). Notably, a similar effect was observed on myeloid cells after treatment with rapamycin (**Figure 9K**). Celecoxib and rapamycin did not affect CD3^+^CD8^+^ T-cell numbers (**Supplemental Figure 10B**). Other lymphocyte populations including CD3^+^CD4^+^ T-cells were only modestly increased in the mutant skin at this stage, and more prominently reduced by rapamycin compared to celecoxib (**Supplemental Figure 10B**).

Collectively, our data show that inhibition of CSF1R or COX-2 limit *Pik3ca*-driven lymphangiogenesis, suggesting that anti-inflammatory therapy provides a potential therapeutic approach for the treatment of LM. Our results also suggest that in addition to inhibiting the downstream PI3K target mTOR in LECs, the immune suppressive effect of rapamycin affecting both myeloid and lymphoid cells may contribute to its therapeutic benefit in the treatment of LM.

## Discussion

Using a mouse model of oncogenic PI3K-driven venous and lymphatic malformations, we characterized endothelial subtype-specific responses to the common causative *Pik3ca^H1047R^* mutation. We uncover a new immunoregulatory subtype of dermal lymphatic capillary endothelial cells, iLECs, that selectively expands in the *Pik3ca^H1047R^*-driven LM. Increased expression of pro-inflammatory factors by iLECs and associated recruitment of myeloid cells in turn support pathological lymphangiogenesis that is inhibited by targeting of the associated immune response.

*PIK3CA* mutations specifically cause vascular malformations in veins and lymphatic vessels. The basis of such vessel type-restricted disease manifestation is unknown. We found that activation of oncogenic p110α-PI3K signaling in the embryonic or postnatal vasculature triggered distinct cellular responses in different vessel types, characterized by vessel sprouting (lymphatic vessels), localized dilation (blood capillaries and veins), or no apparent effect (arteries). Why the same oncogenic stimulus induces different responses in these cells remains unclear, but likely both EC-autonomous and non-autonomous mechanisms play a role.

Modelling of *Pik3ca*-driven venous malformations in the mouse retina recently uncovered that active angiogenesis is required for vascular overgrowth (41), similar to what has been reported in other types of vascular malformations (42–45). Here, we observed a similar vascular response to *Pik3ca^H1047R^* expression in growing embryonic as well as quiescent postnatal dermal blood and lymphatic vessels. Tissue-specific differences in the availability of growth factors that can synergize with the oncogenic p110α-PI3K to enhance downstream signaling (16) may provide a potential explanation for the apparently different vascular responses in the postnatal retina and skin.

In agreement with previous observations (7, 16, 41), proliferation was a common early response of venous ECs and LECs to oncogenic *Pik3ca*. However, proliferation was sustained selectively in LECs of advanced lesions. The first weeks of active lymphatic, but not blood vessel overgrowth was associated with an increase in F4/80^+^ macrophages and Ly6C^+^ proinflammatory monocytes. The latter is particularly noteworthy, given the low abundance of Ly6C^+^ monocytes in healthy tissues and their recruitment from the blood to the site of inflammation, where they differentiate into proinflammatory macrophages (46, 47). Chemokines and cytokines associated with the recruitment of Ly6C^+^ monocytes (CCL2/MCP1) (47) and their proinflammatory signaling (IL1ß, IL6 and TNFα) (48) were also increased. Importantly, macrophages are a major source of pro-lymphangiogenic factors including VEGF-C (22, 23), which was previously shown to be required for the growth of *Pik3ca*-driven lymphatic malformations in mice (16). In support of a role of macrophages in LM growth, we found that blockade of their expansion and differentiation using anti-CSF1R antibody inhibited *Pik3ca*-driven lymphatic overgrowth. Whether macrophages are the only source of VEGF-C in LM remains to be shown.

LECs have been shown to express certain chemokines and cytokines, but their role in paracrine activation of myeloid cells has not been well characterized. Using single cell transcriptomics, we identified a dermal LEC hierarchy that recapitulates the lymphatic vascular architecture of collecting vessels and capillaries *in vivo* (**Figure 10**). This analysis additionally identified a previously unknown and molecularly distinct *Ptx3^high^* population within dermal lymphatic capillary terminals as a putative immunoregulatory LEC subtype, termed iLEC. The iLEC population shared features of a distinct *Ptx3^+^* lymph node LEC subset (49), and was characterized by high expression of transcripts encoding key regulators of innate and adaptive immune responses (*Ptx3, Mrc1, Ackr2*). We observed selective expansion and proliferation of the *Ptx3^high^* iLECs in the *Pik3ca^H1047R^* mice, and high lymphatic endothelial expression of PTX3 in *PIK3CA^H1047R^*-driven lymphatic malformation in humans. Additional upregulation of pro-inflammatory genes, including *Ccl2*, in iLECs suggested a role in paracrine immune activation, and as the pathological cell population in LM.

**Figure 10.**
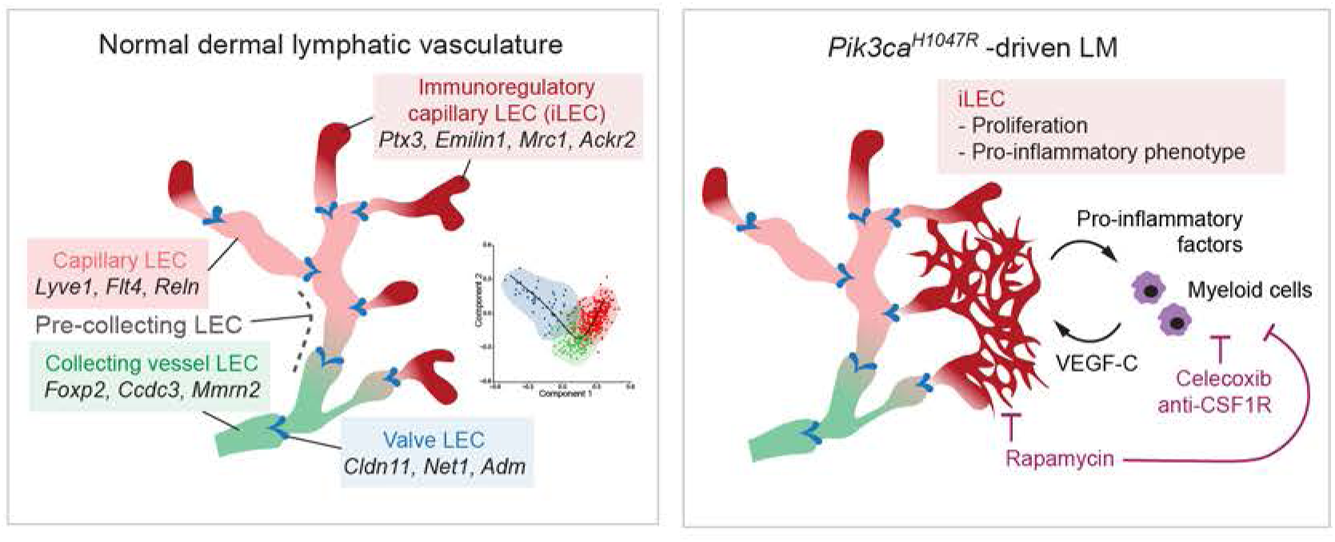
Proposed model of paracrine LEC-immune cell interactions as a driver of lymphatic malformations. Schematic illustration showing distinct dermal LEC subtypes within lymphatic vessel hierarchy (on the left), and pathological changes driven by oncogenic *Pik3ca^H1047R^* (on the right). *Pik3ca^H1047R^* drives expansion and activation of a pro-inflammatory transcriptome in iLECs to induce recruitment and activation of myeloid cells that in turn promote pathological lymphangiogenesis by secretion of pro-lymphangiogenic factors. Celecoxib and anti-CSF1R therapy inhibit lymphangiogenesis by reducing myeloid cell numbers.

Monocyte-derived macrophages are associated with the formation of lymphoid aggregates and tertiary lymphoid organs (TLOs) in chronic inflammation (36), that are frequently found in patients with LMs (37). Targeting of a broader immune response, e.g. by inhibiting the production of prostaglandin, may thus be required for efficient treatment of advanced lesions. We found that systemic COX-2 inhibition using celecoxib limited *Pik3ca^H1047R^*-driven lymphangiogenesis in mice. Inhibition of macrophage recruitment (our study and (38, 39)) or VEGF-C/D production (39, 50, 51) by celecoxib may account for its anti-lymphangiogenic effects. However, other mechanisms including its direct effects on the immunoregulatory functions of LECs themselves (52) or T cells (40) cannot be excluded. Notably, one patient with intractable progressively growing cervical LM was successfully treated with celecoxib (53). Although not curative, celecoxib or other anti-inflammatory treatments could thus provide a clinical benefit in patients with progressive LM.

The mTOR inhibitor rapamycin has provided beneficial effects in clinical trials for VM and LM treatment (1, 2). Because rapamycin induces cell cycle arrest, sustained proliferation of LECs in advanced LM lesions may make them exquisitely sensitive to rapamycin. Rapamycin can exert additional inhibitory effects on LM growth by reducing VEGFR3 levels and thereby upstream lymphangiogenic growth factor signaling in LECs (54, 55). Interestingly, VEGFR3 downregulation is also observed in *Pik3ca* deleted mice and upon treatment with other PI3K pathway inhibitors, the p110α-specific BYL719 (Alpelisib) and a dual PI3K/mTor inhibitor Dactolisib (our unpublished observations). Our results further indicate that rapamycin affects the recruitment of both myeloid and lymphoid cells, which may additionally contribute to its therapeutic benefit in the treatment of LM. Given the multiple effects of PI3K inhibition on LM pathology, as data from clinical trials become available, it will be of interest to investigate whether rapamycin or other PI3K pathway inhibitors provide a better therapeutic outcome in the treatment of *PIK3CA*-driven LMs compared to VMs.

The key advantage of the models utilized in our study is that the inducible, EC-subtype-specific and locally limited activation of *Pik3ca^H1047R^* expression in the dermal vasculature of the ear allows highly reproducible and robust induction of lesions while minimizing potentially life-threatening effects in the internal organs, thereby allowing an extended observation period compared to systemic models. Refined models that combine low-frequency of induction of recombination with tracking of mutant *Pik3ca^H1047R^*-expressing ECs to induce single isolated vascular lesions will be useful to study clonal dynamics and to allow further extending the time period of lesion formation in future studies. This is likely required to fully recapitulate the chronic inflammation observed in human patients to allow assessing the therapeutic benefit of targeting the immune response in the treatment of advanced lesions.

Taken together, our study reveals a new immunoregulatory subtype of dermal lymphatic capillary endothelial cells, iLECs, as a pathological cell population in LM that contributes to paracrine immune activation that in turn supports pathological lymphangiogenesis. Apart from identifying the immune response as a therapeutic target for the treatment of LM, our findings have implications for understanding the role of lymphatic vessels as upstream orchestrators of the immune response in other inflammatory conditions.

## Methods

### Mouse lines and treatments

*R26-LSL-Pik3ca^H1047R^* (56), *R26-mTmG* (57) and *tdTom* (58) mice were crossed with the *Cdh5-CreER^T2^* (21), *Vegfr3-CreER^T2^* (20) or *Vegfr1-CreER^T2^* (generated in this study, see supplemental material). Both female and male mice were used for analyses and no differences in the phenotype between the genders were observed. The morning of vaginal plug detection was considered as embryonic day 0 (E0). Cre-mediated recombination was induced in embryos by intraperitoneal injection of 0.5 mg (*Cdh5-CreER^T2^*) or 2 mg (*Vegfr1-CreER^T2^, Vegfr3-CreER^T2^*) of 4-hydroxytamoxifen (4-OHT, H7904, Sigma), dissolved in peanut oil (10 mg/ml), to pregnant females at E10 (*Vegfr1-CreER^T2^*), E11 (*Cdh5-CreER^T2^, Vegfr3-CreER^T2^*) or at E14 (*Cdh5-CreER^T2^*). For postnatal induction, 25 μg (*Cdh5-CreER^T2^*) or 50 μg (*Vegfr3-CreER^T2^* and *Vegfr1-CreER^T2^*) of 4-OHT dissolved in acetone (10 mg/ml) was administered topically to the dorsal side of each ear of 3-week-old mice. Littermate controls were included in each experiment, which were either Cre^−^ mice treated with 4-OHT (*Cdh5-CreER^T2^*, *Vegfr1-CreER^T2^*), or Cre^+^ mice treated with the vehicle (*Vegfr3-CreER^T2^*). 4-OHT-treated mice carrying the different Cre alleles in combination with the *R26-mTmG* reporter showed normal vasculature, excluding Cre-induced toxicity (59) as the cause of the phenotypes in the *Pik3ca*-expressing mice. Celecoxib (LC Laboratories, C-1502) was dissolved first in DMSO (400 mg/ml) followed by mixing 1:20 with oil (final concentration of 20 mg/ml), and administered by oral gavage every second day at the dose of 100 mg/kg. InVivoMAb anti-mouse CSF1R antibody (BioXCell, clone AFS98, BE0213) was diluted in PBS and administered by intraperitoneal injection (30 mg/kg) twice per week. Control mice were administered with the vehicle.

### Generation of *Vegfr1-CreER^T2^* mice

To generate transgenic mice expressing the tamoxifen inducible *CreER^T2^* under the control of the *Flt1* (*Vegfr1*) promoter, a BAC (B6Ng01-247E8) carrying the mouse *Vegfr1* gene (RefSeq NM_010228.3) was obtained from the mouse B6Ng01 library. The BAC vector was engineered to replace the coding sequence in exon 1 as well as the splice donor site at the junction between exon 1 and intron 1 (50 bp) with a cassette containing the open reading frame for *CreER^T2^* and the *Vegfr1* 3’ untranslated region (UTR), such that the endogenous translation initiation codon from the *Vegfr1* gene is used for the expression of the CreER^T2^ protein. A polyadenylation signal (human Growth Hormone polyadenylation signal) was inserted 3’ of the *Vegfr1* 3’UTR (**Supplemental Figure 1A**). The engineered BAC vector was used for pronuclear injection into fertilized C57BL/6NTac oocytes. Two independent founder lines were generated, of which one showed efficient Cre-mediated recombination in the *Vegfr1*-expressing cells and was used for further analyses. Mice were generated by Taconic Biosciences.

Transgenic mice were detected by PCR with primers designed to amplify a 494 base pair region at the junction of 5’ mouse genomic region and the *CreER^T2^* open reading frame in the transgene (forward 19966_7: CACTTCAGCGAGGTCCTTGAG5 and reverse 19966_112: CATCTTCAGGTTCTGCGGG). Additional control primers (forward 11767_3: GGGGCAATCAATTGAGGG and reverse 11767_4: CAACCTCTGCTTGGTTCTGG) were included in the reaction to amplify a 333 bp fragment. Standard amplification reactions (25 μl) were prepared using 0.4 μM of each primer and 0.2 mM dNTPs. After initial denaturation at 95°C for 5 minutes, reactions were subjected to 35 cycles of 95°C (30 s), 60°C (30 s), and 72°C (60 s). Reactions were incubated for a final elongation at 72°C for 10 minutes. PCR products were separated on a 2% agarose gel containing Sybr Safe.

### Antibodies

The details of primary antibodies used for immunofluorescence of whole mount tissues and flow cytometry are provided in **Supplemental Table 2**. Secondary antibodies conjugated to AF488, AF594, AF647 or Cy3 were obtained from Jackson ImmunoResearch and used in 1:300 dilution (**Supplemental Table 2**).

### Whole-mount immunofluorescence

Tissues (skin, diaphragm or intestine wall) were fixed in 4% paraformaldehyde for 2 h at RT and permeabilized in 0.3% Triton X-100 in PBS (PBST) for 10 min. After blocking in PBST with 3% milk for at least 1.5 h, the tissues were incubated with primary antibodies at 4°C overnight in blocking buffer, followed by PBST washing and incubation with fluorescence-conjugated secondary antibodies for 2 h at RT. After washes in PBST the samples were mounted in Mowiol. PTX3 and CCL2 staining was amplified by Tyramide Signal Amplification kit (TSA^TM^, NEN^TM^ Life Science Products). Tissue was first blocked with TNB (Tris-NaCl-blocking buffer), prepared according to the TSA kit instructions. For CCL2 staining the rest of the protocol was done according to TSA kit instructions with PBST used as a permeabilization reagent. PTX3 staining and washing of the tissue was done in PBS in the absence of permeabilization agent to allow visualization of extracellular proteins.

### Edu Click-IT Kit assay

DNA synthesis in proliferating cells was detected using the Click-iT™ EdU Cell Proliferation Kit for Imaging (Thermo Fisher Scientific). Mice received an intraperitoneal injection with 25 mg/kg of Edu 16 hours prior to harvesting the ears for analysis. After whole-mount immunofluorescence staining (specified above, except for PROX1 staining that was performed after the Click-IT assay), the tissues were washed extensively with PBST and stained for EdU according to the manufacturer’s instructions. Briefly, tissues were incubated in the Click-IT Reaction cocktail for 40 min at RT followed by washing in PBST.

### Confocal microscopy and image processing

All confocal images were acquired using a Leica SP8 or a Leica STELLARIS 5 confocal microscope equipped with a white light laser and 10×/0.45 C-Apochromat (HC PL APO CS2), 20×/0.75 (HC PL APO CS2), 25×/0.95 (HC FLUOTAR L VISIR), or or 63x/1.20 (HC PL APO) objective. The images were obtained at room temperature using Leica LAS X software. All images were processed using Fiji Image J (NIH) software. Each image represents maximum intensity projection of a Z-stack (capturing the entire lymphatic vascular layer, or the whole tissue Z-stack when imaging immune populations) of single tiles or multiple tile scan images. The ear tile scans in Supplemental Figure 2 were obtained using DMI8 Leica fluorescence microscope (Supplemental Figure 2A) or Leica Thunder Imaging System (Supplemental Figure 2B). The close-up images in Figure 2, D and E, and Figure 3, A and C, Figure 5F, Figure 6I, Figure 8D and Supplemental Figure 5C and 7, A and B were additionally deconvolved using Huygens Essential (v 19.04) (Scientific Volume Imaging) with theoretical point spread function and automatic background estimation. Details of image processing and quantification are provided in supplemental material.

### FACS analysis

FACS analysis of Ki67^+^ ECs and CD45^+^CD11b^+^F4/80^+^ cells was done as previously described (*16*). For FACS analysis of innate and adaptive immune cells, the ear skin was dissected, cut into pieces, and digested in Liberase TL (100 µg/ml, Roche) (for innate immune cells) or Collagen IV (2 mg/ml, Life Technologies) (for adaptive and NK cells) plus DNase I (0.5 mg/ml, Roche) in PBS with 0.2% FBS at 37 °C for 2 h at 700 rpm. The cell lysate was filtered through 50 µm filters (CellTricks) and washed with FACS buffer. The cells were first incubated with rat anti-mouse CD16/32 antibody (eBioscience) to block Fc receptor binding. Cell suspensions were stained for antibodies targeting different immune populations (**Supplemental Table 2**). Dead cells were labelled using LIVE/DEAD™ Fixable Near-IR Dead Cell Stain Kit (Life Technology). The analysis was performed on BD LSRFortessa^TM^

Cell Analyser (BD Biosciences). All data was processed using FlowJo software version 10.5.0 (TreeStar). Gating of Ki67^+^ ECs and CD45^+^CD11b^+^F4/80^+^ was done as previously described (*16*). Gating scheme for innate and adaptive immune cells is shown in **Supplemental Figure 11**. Relative cell frequency of the subtypes of immune populations are presented as fold change (in % of all live cells) relative to the average of the control in each experiment.

### Single cell transcriptomics

Dermal LECs and BECs were FACS-sorted from the ear skin of 4-OHT-treated 5-week-old *Pik3ca^H1047R^;Cdh5-CreER^T2^* (*n*=5) and Cre^−^ littermate control mice (*n*=2) of mixed genders. At this stage, robust vascular overgrowth, high LEC proliferation and immune cell infiltration were observed in the mutants. One wild type C57BL/6J mouse, not treated with 4-OHT, was also included, to exclude possible effects of the solvent and 4-OHT included in the treatment. Ear skin was first digested in 5 mg/ml Collagenase II in PBS supplemented with 0.2 mg/ml DNaseI and 0.2% FBS, followed by filtering through 50 µm filters (CellTricks). Fc-receptors were blocked with rat anti-mouse CD16/32 antibody (eBioscience), followed by staining using Podoplanin-APC and CD31-Pe-Cy7 antibodies. Dump channel included erythrocytes and immune cells (labeled using CD45-eF450, CD11b-eF450 and Ter119-eF450) and dead cells (SYTOX Blue dead stain (Life Technology)). The sorting into 384-well plates was performed with a 100 µm nozzle on BD FACS AriaIII CellSorter (BD BioScience Flow Cytometry System equipped with four lasers: 405, 488, 561, 633 nm). Smart-Seq2 library preparation and sequencing were performed as described previously (60). Key quality metrics are listed in **Supplemental Table 3**.

### scRNAseq data processing

The single cell sequence data were aligned to the mouse reference genome GRCm38 with tophat (version 2.1.1) (61) and *Gallus gallus PIK3CA* sequence (NCBI nucleotide sequence id: NM_001004410). The latter was used to identify the *Pik3ca^H1047R^* transcript in the transgenic mice, encoded by *Gallus gallus Pik3ca*, which is highly homologous (83%) to the mouse *Pik3ca* (NM_008839.3). The corresponding protein sequence of *Gallus gallus* (NP_001004410) is 96% identical to mouse (NP_032865.2), and oncogenic in mammalian cells (56, 62). Duplicated reads were filtered out using samtools software (version 0.1.18). The gene counts were summarized using featureCounts function from the Subread package (version 1.4.6-p5) (63). Further downstream analysis of raw expression data was performed in RStudio (desktop version 2021.09.2+382) using R versions 3.5.1 and 4.0.3 (64, 65). The following quality control steps were performed: (1) genes expressed by less than 3 cells were removed; (2) cells with fewer than 200 genes or 50,000 reads counts were not further considered; (3) cells in which over 10% reads derived from the mitochondrial genome were removed. After removing cells with poor transcriptome quality, the data was processed in Seurat package (version: 3.1.1 and 4.0.1) (28, 66) for normalization using *LogNormalize* function, graph-based clustering analysis (Louvain method), non-linear dimensional reduction (UMAP) and differentially expressed gene (DEG) detection for identification of cluster markers (Wilcoxon Rank Sum test, marker genes selected by p value with Bonferroni correction less than 0.05 and logarithmic fold change greater than 1). The data was further processed in two steps in which a LEC population was extracted from the control and mutant datasets individually after removal of contaminating cells identified as epithelial cells/keratinocytes, fibroblasts, mural cells and immune cells based on DEG analysis. In the second step, LECs from the control and mutant mice were integrated, and additional small clusters with neuronal, immune cell and fibroblast identities were removed. Batch correction and data integration was performed using canonical correlation analysis (CCA) method for control data, and Harmony package (version: 1.0) (67) for mutant and integrated control/mutant data.

### Trajectory Inference analysis

Trajectory inference (TI) analysis was performed using *SCORPIUS* (v.1.0.5, (68)) for the control dataset (Figure 5C) and *SLINGSHOT (v.2.2.0)* (69) for the combined dataset (Figure 6D) in which the respective algorithm constructed the topology of dynamic process as a linear trajectory and mapped the cells along this trajectory curve. In the process of constructing the trajectory using *SCORPIUS*, we used *k*=3 and other parameters as default. To select the markers to be able to significantly predict the order of the cells along the trajectory, we applied Random Forest regression model (RF) (70) using *randomForest* package (v. 4.6-14). For each tree, RF model was trained and prediction error rate was measured by using out-of-bag (OOB) data; in total 1000 trees were generated. A variable importance (VI) score, Mean Decrease Accuracy, was calculated for all variables/genes, and genes were ranked in descending order by VI score. Using *SLINGSHOT* two lineages/branches were obtained by constructing minimum spanning trees on clusters in an unsupervised manner using default parameters without forcing a cluster of origin or leaf node.

### GO analysis

For Gene Ontology (GO) enrichment analysis on the biological processes domain, the hypergeometric test in the *GOstats* package (version: 2.56.0) was applied. The analysis was restricted to gene sets containing 5-1000 genes. Significant pathways were filtered by applying *p* value <0.05 and gene count/term >10. Further selection of relevant GO terms was based on sorting terms on their OddsRatios (the ratio of a GO term in the differently expressed genes list to the occurrence of this GO term in a universal gene list, obtained from org.Mm.eg.db (version 3.13.0)). In addition, GO term results were screened for enrichment of terms relate to immune regulation. Selected top pathways were visualized using ggplot2 (version: 3.3.5).

### RNA extraction and real-time qPCR

Total RNA was extracted from LECs, BECs, or FACS-bulk-sorted as previously described for the scRNAseq analysis, as well from CD45^+^CD11b^+^Ly6G^−^F480^+^ macrophages and CD45^+^CD11b^+^Ly6G^−^F480^+^Ly6C^+^ monocytes, by using RNeasy Micro Kit (QIAGEN), according to the manufacturer’s instructions with additional DNaseI treatment. The cDNA was synthesized using Superscript VILO Master Mix (Invitrogen) and TaqMan™ PreAmp Master Mix was used for preamplification with subsequent real-time qPCR on StepOne Plus Real-Time PCR system (Applied Biosystems) using TaqMan gene expression Master Mix (Applied Biosystems). The following TaqMan Assays were used: *Hprt* (Mm01545399_m1), *Mki67* (Mm01278617_m1) and *Vegfc* (Mm00437310_m1). Relative gene expression levels were calculated using the comparative CT method with *Hprt* as reference gene.

### VEGF-C ELISA

Mouse Vascular Endothelial Cell Growth Factor C (VEGF-C) ELISA Kit from CUSABIO (CSB-E07361m) was used for detection of VEGF-C protein concentration. Tissue homogenates were prepared according to the instructions of the kit using freeze-thaw cycles for disrupting the cell membrane. For normalization, the protein concentration of the sample was measured using Thermo Scientific™ Pierce™ BCA Protein Assay Kit. Subsequent steps were performed according to the manufacturer’s instructions.

### Multiplex ELISA

Meso Scale Diagnostics (MSD) Multiplex ELISA using MSD Multi-Spot Assay System was used with two different V-PLEX MSD cytokine assays for detection of total 19 proteins, divided into two panels – Proinflammatory Panel 1 (including IFN-γ, IL-1β, IL-2, IL-4, IL-5, IL-6, CXCL1 (KC/GRO), IL-10, IL-12p70, TNF-α) and Cytokine Panel 1 (IL-9, MCP-1, IL-33, IL-27p28/IL-30, IL-15, IL-17A, MIP-1a, IP-10, MIP-2). Ear skin lysates were prepared by homogenizing the tissue with Homogenizer TissueRuptor II (QIAGEN) in TBS + 1% Triton-X100 + Protease and Phosphatase Inhibitor Cocktail (Sigma). Protein concentration was measured and adjusted to 100 mg/ml before adding the samples to 96-well plate-based multiplex assay plate (provided by Meso Scale Discovery Kit) containing detection antibodies, conjugated with electrochemiluminescent labels (MSD SULFO-TAG™). Subsequent steps were performed according to the manufacturer’s instructions. Samples which gave a readout below the manufacturer’s control threshold were excluded.

### Analysis of human tissue

Human biopsy material was obtained from Xarxa de Bancs de Tumors de Catalunya (XBTC) biobank, approved by PIC-96-16. Histology was examined by a pathologist expert in vascular anomalies and summarized in **Supplemental Table 1**. PTX3 staining of paraffin sections was amplified by Tyramide Signal Amplification kit (TSA^TM^, NEN^TM^ Life Science Products). Tissue was blocked with TNB (Tris-NaCl-blocking buffer), prepared according to the TSA kit instructions.

### Statistics

Graphpad Prism 7-9 was used for graphic representation and statistical analysis of the data. Data between two groups were compared with unpaired Student’s t-test. Differences were considered statistically significant when *p*<0.05 and indicated on the graphs with star symbols: **** *p*<0.0001, *** *p*<0.001, ** *p*<0.01, * *p*<0.05. For identifying cluster markers from scRNA-seq data, Wilcoxon Rank Sum test was performed. Marker genes were selected by *p* value with Bonferroni correction less than 0.05 and logarithmic fold change greater than 1.

### Study approval

Experimental procedures on mice were approved by the Uppsala Animal Experiment Ethics Board (permit numbers 130/15, 5.8.18-06383/2020) and performed in compliance with all relevant Swedish regulations. Human biopsy material was obtained from Xarxa de Bancs de Tumors de Catalunya (XBTC) biobank, approved by PIC-96-16. Analysis of human biopsy material was approved by Swedish Ethical Review Authority (Etikprövningsmyndigheten, permit number: Dnr 2020-00987).

### Data availability

The data that support the findings of this study are available from the corresponding author upon reasonable request. The single cell raw sequencing data and processed counts tables, as well as R files containing raw counts and metadata for Figure 5B and Figure 6, A and D have been deposited in GEO (Gene Expression Omnibus) repository and will be made available when the manuscript has been accepted for publication.

## Author contributions

MP, MK, SS, IMC and TM conceived and designed the study. MP, MK, SS, IMC, HO and BJ designed and performed experiments. MK and YS performed scRNAseq data analysis. MP, MK, SS, IMC and TM analyzed and interpreted data. SDC, EB and MG provided human samples. MV and CB provided scRNAseq resources. TM supervised the project. MP and TM wrote the manuscript. All authors discussed the results and commented on the manuscript.

## Acknowledgements

We thank Dieter Saur (Technische Universität München) for the *R26-LSL-Pik3ca^H1047R^* mice, Sagrario Ortega (CNIO, Madrid) for the *Vegfr3*-*CreER^T2^* mice and Ralf Adams (Max Planck Institute for Molecular Biomedicine, Münster) for the *Cdh5*-*CreER^T2^* mice. We also thank the BioVis facility (Uppsala University) for flow cytometer usage and support, the Single cell core Facility at Flemingsberg campus (SICOF), Karolinska Institute, for single cell sequencing services, Maria Globisch for advice and help with the Multiplex ELISA, and Sofie Lunell Segerqvist and Aissatu Mami Camara for technical assistance. The computations of single cell RNA sequencing data were enabled by resources provided by the Swedish National Infrastructure for Computing (SNIC) at Uppsala Multidisciplinary Center for Advanced Computational Science (UPPMAX) partially funded by the Swedish Research Council through grant agreement no. 2018-05973, under projects SNIC 2018/8-62 and SNIC 2020/16-159. Prasoon Agarwal at LUNARC is acknowledged for assistance with the analysis of scRNAseq data.

This work was supported by grants from Knut and Alice Wallenberg Foundation (2018.0218 (TM) and 2020.0057 (TM, CB)), the Swedish Research Council (2020-0269) (TM), Göran Gustafsson foundation (TM), the Swedish Cancer Society (19 0220 Pj, 19 0219 Us) (TM), the European Research Council (ERC-2014-CoG-646849) (TM), and the European Union’s Horizon 2020 research and innovation programme under the Marie Skłodowska-Curie grant agreement No 814316 (TM, MK), La Caixa Foundation (HR18-00120) (MG, EB), and la Caixa Banking Foundation (LCF/BQ/PR20/11770002) (SDC). SS was supported by a research fellowship from the Deutsche Forschungsgemeinschaft (STR 1538/1-1) and a non-stipendiary long-term fellowship from the European Molecular Biology Organization (ALTF 86-2017). SDC is a recipient of a fellowship from the European Union’s Horizon 2020 Research and Innovation Programme under the Marie Sklodowska-Curie grant agreement No 749731.

The authors have no conflicting financial interests.

## Supplemental material

### Supplemental Tables

**Supplemental Table 1.** Clinical features of patients with LM driven by *PIK3CA^H1047R^* mutation.

| ID | Lesion type | Gender | Age at biopsy | Location | Diagnosis |
| --- | --- | --- | --- | --- | --- |
| Case1<br>VM168 | Lymphatic Malformation (microcystic) | M | 17 y | Skin (leg) | Epidermis showing acanthosis with hyperkeratosis and papillomatosis; Presence of ectatic and congestive dilated vessels containing erythrocytes in superficial dermis; Mild chronic perivascular inflammation. Endothelium PECAM1<sup>+</sup>, GLUT1<sup>-</sup>. |
| Case2<br>VM130 | Mixed Veno-Lymphatic Malformation | F | 13 y | Nose | Fibrofatty tissue including some skeletal muscle fibers; Presence of vessels with irregular morphology, some of them with muscular wall and evidence of lymphoid aggregates. Endothelium PECAM1<sup>+</sup>, PDPN<sup>+/-</sup>, GLUT1<sup>-</sup>. |
| Case3<br>VM97 | Lymphatic Malformation (microcystic) | F | 7 y | Tongue | Tissue fragments covered by squamous epithelium with hyperplastic changes. Subepithelial stroma contains dilated vascular structures with serous material, lined by endothelium without atypia. Accompanying chronic inflammation. Endothelium PECAM1<sup>+</sup>, PDPN<sup>+</sup>, GLUT1<sup>-</sup>. |

**Supplemental Table 2.** List of antibodies.

| **Primary antibody: Immunostaining** | **Company and catalog number** | **Dilution** |
| --- | --- | --- |
| chicken anti-GFP | Abcam, ab13970 | 1/300 |
| goat anti-mouse NRP2 | R&D Systems, AF567 | 1/200 |
| goat anti-mouse VEGFR3 | R&D Systems, AF743 | 1/100 |
| goat anti-human PROX1 | R&D Systems, AF2727 | 1/100 |
| hamster anti-mouse PDPN-AF488 | eBioscience, 53-5381-82 | 1/50 |
| hamster anti-mouse PDPN (clone 8.1.1) | Developmental Studies Hybridoma Bank, 8.1.1-a | 1/200 |
| rabbit anti-mouse LYVE1 | Reliatech, 103-PA50AG | 1/500 |
| rabbit anti-mouse Ki67 (clone SP6) | ThermoFischer, MA5-14520 | 1/100 |
| rat anti-mouse PECAM1 | Becton Dickinson, 553370 | 1/200 |
| rat anti-mouse EMCN | Santa Cruz Biotechnology, sc-65495 | 1/200 |
| rat anti-mouse CD45 | Abcam, ab25386 | 1/100 |
| rat anti-mouse CD45-FITC | eBioscience, 11-0451 | 1/50 |
| mouse anti-Actin α-Smooth muscle-Cy3 | Sigma, clone 1A4, C6198 | 1/500 |
| rat anti-mouse LYVE1 | R&D Systems, MAB2125 | 1/200 |
| rabbit anti-mouse Collagen IV | Bio-Rad, AbD Serotec, 2150-1470 | 1/1000 |
| rabbit anti-mouse PTX3 | Thermo Fischer, PA5-38595 | 1/100 |
| rat anti-mouse Ly-6C (HK1.4, APC) | BioLegend, 128015 | 1/50 |
| rat anti-mouse F4/80 (clone Cl:A3-1) | BioRad, MCA497GA | 1/100 |
| hamster anti-mouse PECAM1 (clone 2H8) | Thermo Fischer, MA3105 | 1/300 |
| goat anti-mouse-CCL2 | R&D Systems, AF-479 | 1/100 |
| rabbit anti-human PTX3 | Sigma, HPA069320 | 1/100 |
| mouse anti-human Podoplanin-AF594, clone D2-40 | BioLegend, 916607 | 1/100 |

| **Secondary antibody: Immunostaining** | **Company and catalog number** | **Dilution** |
| --- | --- | --- |
| donkey anti-rat IgG-Cy3 | Jackson ImmunoResearch (JIR), 712-165-153 | 1/300 |
| donkey anti-rat IgG-AF488 | JIR, 712-545-153 | 1/300 |
| donkey anti-rat IgG-AF647 | JIR, 712-605-153 | 1/300 |
| donkey anti-rat-Biotin SP | JIR, 712-065-153 | 1/300 |
| goat anti-Syrian hamster-A5F94 | JIR, 107-585-142 | 1/300 |
| donkey anti-goat-AF647 | JIR, 705-605-147 | 1/300 |
| donkey anti-goat-AF594 | JIR, 705-585-147 | 1/300 |
| donkey anti-rabbit-AF488 | JIR, 711-545-152 | 1/300 |
| donkey anti-rabbit-A647 | JIR, 711-605-152 | 1/300 |
| donkey anti-rabbit-Biotin SP | JIR, 711-065-152 | 1/300 |
| rabbit-anti-hamster-Cy3 | JIR, 307-165-003 | 1/300 |
| donkey anti-rabbit-Cy3 | JIR, 711-166-152 | 1/300 |

| **Primary antibody: FACS** | **Company and catalog number** | **Dilution** |
| --- | --- | --- |
| rat anti-mouse CD16/CD32 | eBioscience, 14-0161 | 1/100 |
| rat anti-mouse PECAM1/CD31 (390, PE-Cyanine7) | eBioscience, 25-0311 | 1/300 |
| hamster anti-mouse PDPN (8.1.1, APC) | eBioscience, 12-7410 | 1/100 |
| hamster anti-mouse PDPN (8.1.1, PE) | eBioscience, 12-5381 | 1/300 |
| rat anti-mouse CD11b<br>(M1/70, PerCP-Cyanine5.5) | eBioscience, 45-0112 | 1/50 or<br>1/100 |
| rat anti-mouse CD11b (M1/70, APC) | eBioscience, 17-0112 | 1/100 |
| rat anti-mouse CD11b (M1/70, eF450) | eBioscience, 48-0112 | 1/50 |
| rat anti-mouse CD45<br>(30-F11, PerCP-Cyanine5.5) | eBioscience, 45-0451 | 1/100 |
| rat anti-mouse CD45 (30-F11, PerCP) | BD Pharmingen, 55-7235 | 1/100 |
| rat anti-mouse CD45 (30-F11, V500) | BD Pharmingen, 56-1487 | 1/100 |
| rat anti-mouse CD45 (30-F11, eF450) | eBioscience, 48-0451 | 1/50 |
| rat anti-mouse F4/80 (BM8, FITC) | BioLegend, 123108 | 1/50 |
| rat anti-mouse Ki67 (SolA15, eFluor 660) | eBioscience, 50-5698 | 1/100 |
| rat anti-mouse TER-119 (TER119, eF450) | eBioscience, 48-5921 | 1/100 |
| rat anti-mouse CD4 (RM4-5, APC) | eBioscience, 17-0042 | 1/100 |
| rat anti-mouse CD3 (17A2, FITC) | eBioscience, 100203 | 1/100 |
| rat anti-mouse B220 (RA3-6B2, eF450) | eBioscience, 48-0452 | 1/100 |
| rat anti-mouse CD8 (53-6.7, SB780) | eBioscience, 78-0081 | 1/100 |
| rat anti-mouse NK (PK136, PE/Cy7) | BioLegend, 108713 | 1/100 |
| rat anti-mouse Ly-6C (HK1.4, APC) | BioLegend, 128015 | 1/100 |
| rat anti-mouse Ly-6G (1A8, PE/Cy7) | BioLegend, 127617 | 1/100 |
| hamster anti-mouse CD11c (N418, BV605) | BioLegend, 117333 | 1/100 |

**Supplemental Table 3.** Key quality metrics for dermal EC scRNA-seq data.

| **Sample ID** | **Mouse genotype** | **Total no of reads** | **Median no of reads** | **Reads/ cell** | **Genes/ cell** |
| --- | --- | --- | --- | --- | --- |
| 895s | Ctrl<br>C57BL/6J | 110 103 815 | 277 427 | 286 729 | 6 527 |
| 901s | Ctrl<br>Cre- littermate | 111 465 290 | 280 166 | 290 274 | 5 054 |
| 903s | Ctrl<br>Cre- littermate | 121 842 883 | 346 507 | 317 299 | 4 294 |
| 891_echo | Mutant<br>*Pik3ca*<sup>H1047R</sup>; *Cdh5-CreER*<sup>T2</sup> | 149 568 482 | 381 729 | 389 501 | 6 558 |
| 892s | Mutant<br>*Pik3ca*<sup>H1047R</sup>; *Cdh5-CreER*<sup>T2</sup> | 151 102 969 | 367 961 | 393 497 | 5 943 |
| 323s | Mutant<br>*Pik3ca*<sup>H1047R</sup>; *Cdh5-CreER*<sup>T2</sup> | 105 188 400 | 245 277 | 273 928 | 6 213 |
| 326s | Mutant<br>*Pik3ca*<sup>H1047R</sup>; *Cdh5-CreER*<sup>T2</sup> | 99 738 886 | 254 315 | 259 737 | 4 837 |
| 329s | Mutant<br>*Pik3ca*<sup>H1047R</sup>; *Cdh5-CreER*<sup>T2</sup> | 121 560 970 | 301 207 | 316 565 | 4 877 |
| 330s | Mutant<br>*Pik3ca*<sup>H1047R</sup>; *Cdh5-CreER*<sup>T2</sup> | 99 829 512 | 196 190 | 259 973 | 5 931 |
| 332s | Mutant<br>*Pik3ca*<sup>H1047R</sup>; *Cdh5-CreER*<sup>T2</sup> | 127 499 101 | 292 351 | 332 029 | 5 187 |

**Supplemental Figure 1.**
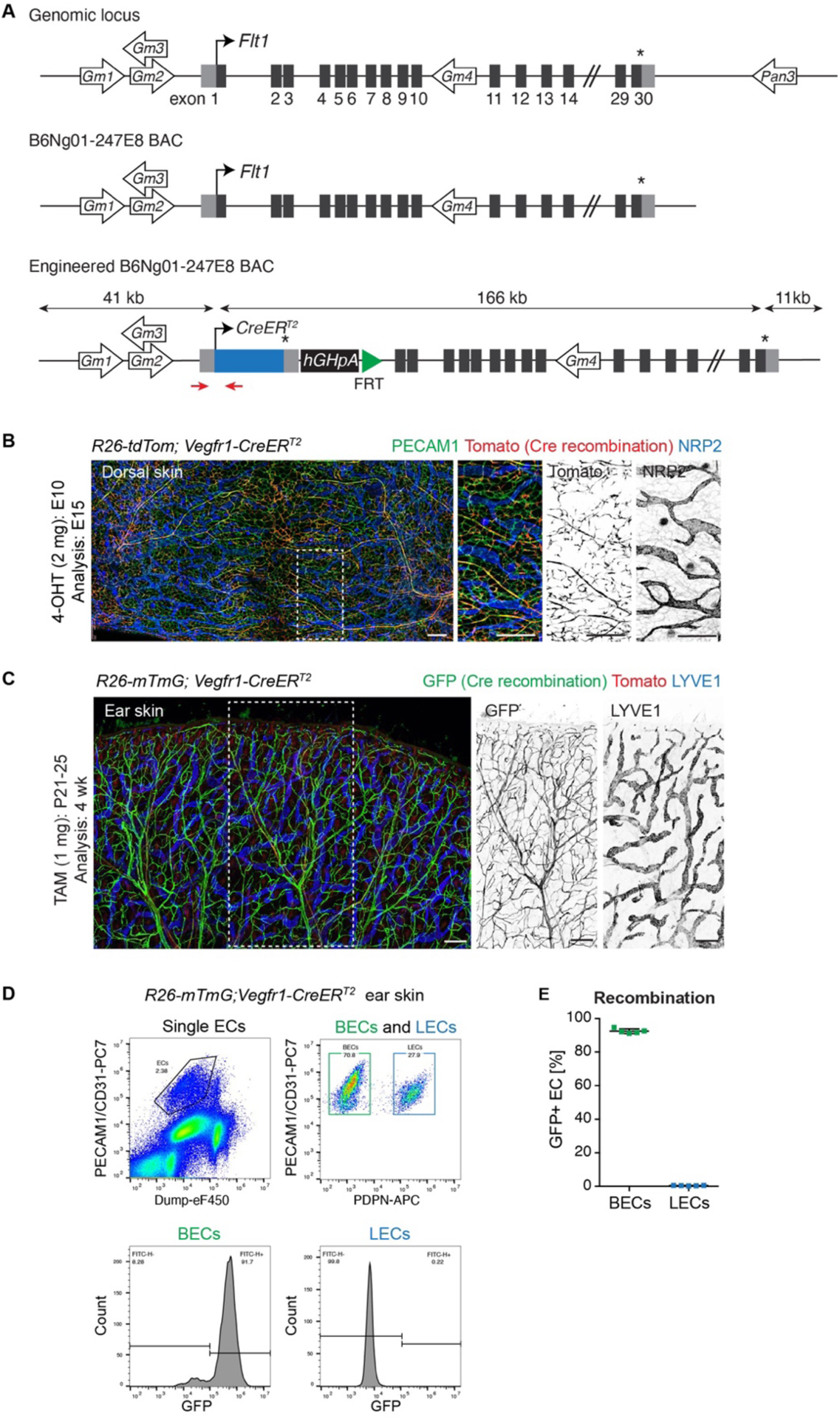
Generation of BAC transgenic *Vegfr1-CreER^T2^* mice. (**A**) Schematic of the genomic locus, a BAC for the *Flt1 (Vegfr1)* gene, and the engineered BAC used for generation of transgenic mice where exon 1 of *Vegfr1* is replaced with the open reading frame for *CreER^T2^*, the *Vegfr1* 3’ UTR and hGH polyadenylation signal. In addition to the *Vegfr1* gene, the engineered BAC also carries the *Gm35592* ncRNA (NCBI gene ID: 102639236), the *Gm35541* ncRNA (NCBI gene ID: 102639166), the *Gm43156* lincRNA (Ensembl gene ID: ENSMUSG00000106845) and the *Gm35383* ncRNA (NCBI Gene ID: 102638948). The location of the genotyping primers are indicated (red arrows). (**B**) Whole-mount immunofluorescence of the skin of E15 *R26-tdTom*;*Vegfr1-CreER^T2^* embryo showing tdTom expression (Cre-mediated recombination) specifically in the PECAM1^+^LYVE1^−^ blood vessels. Boxed area is magnified on the right. 4-OHT was administered at E10. (**C**) Whole mount immunofluorescence of 4-week-old *R26-mTmG*;*Vegfr1-CreER^T2^* ear skin showing GFP expression (Cre-mediated recombination) specifically in the LYVE1^−^ blood vessels. Single channel images are shown for the boxed area. (**D**, **E**) Flow cytometry analysis of dermal ECs from *R26-mTmG*;*Vegfr1-CreER^T2^* ear showing efficient recombination (GFP expression) specifically in the BECs. Data in (E) represent mean (*n*=5 mice) ± s.d. In (C-E), mice received 5 consecutive administrations of 1 mg of tamoxifen at 3 weeks of age and were analyzed at 4 weeks of age. Scale bars: 250 µm (B), 200 µm (C).

**Supplemental Figure 2.**
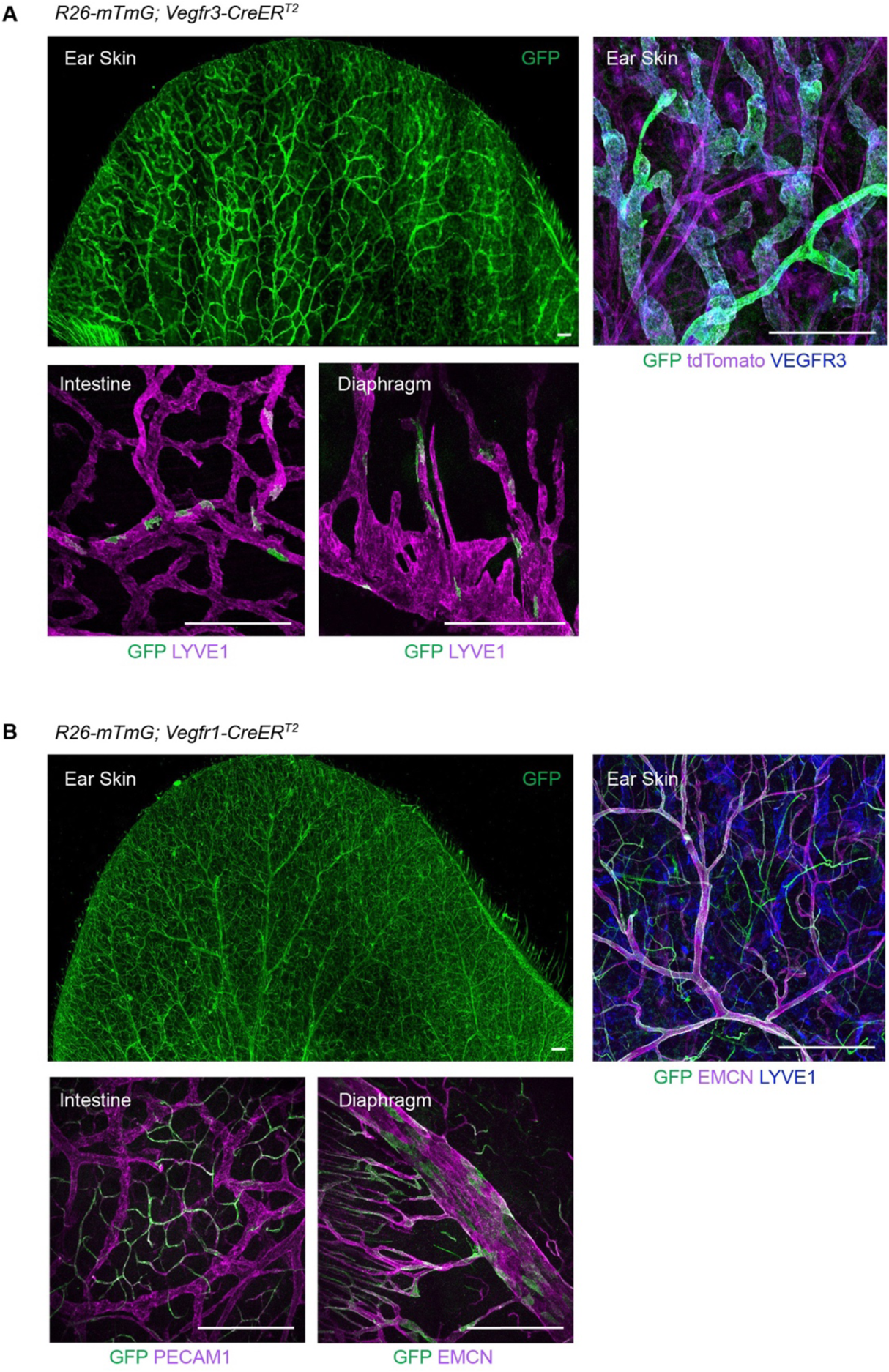
Cre recombination efficiency in the *Vegfr3-CreER^T2^* and *Vegfr1-CreER^T2^* mice after topical 4-OHT administration. GFP expression in a whole-mount ear skin and the internal organs of *R26-mTmG;Vegfr3-CreER^T2^*(**A**) and *R26-mTmG;Vegfr1-CreER^T2^* (**B**) mice after topical application of 4-OHT to the ears. Systemic recombination was observed at a low efficiency in the intestine and diaphragm. Scale bars: 400 µm.

**Supplemental Figure 3.**
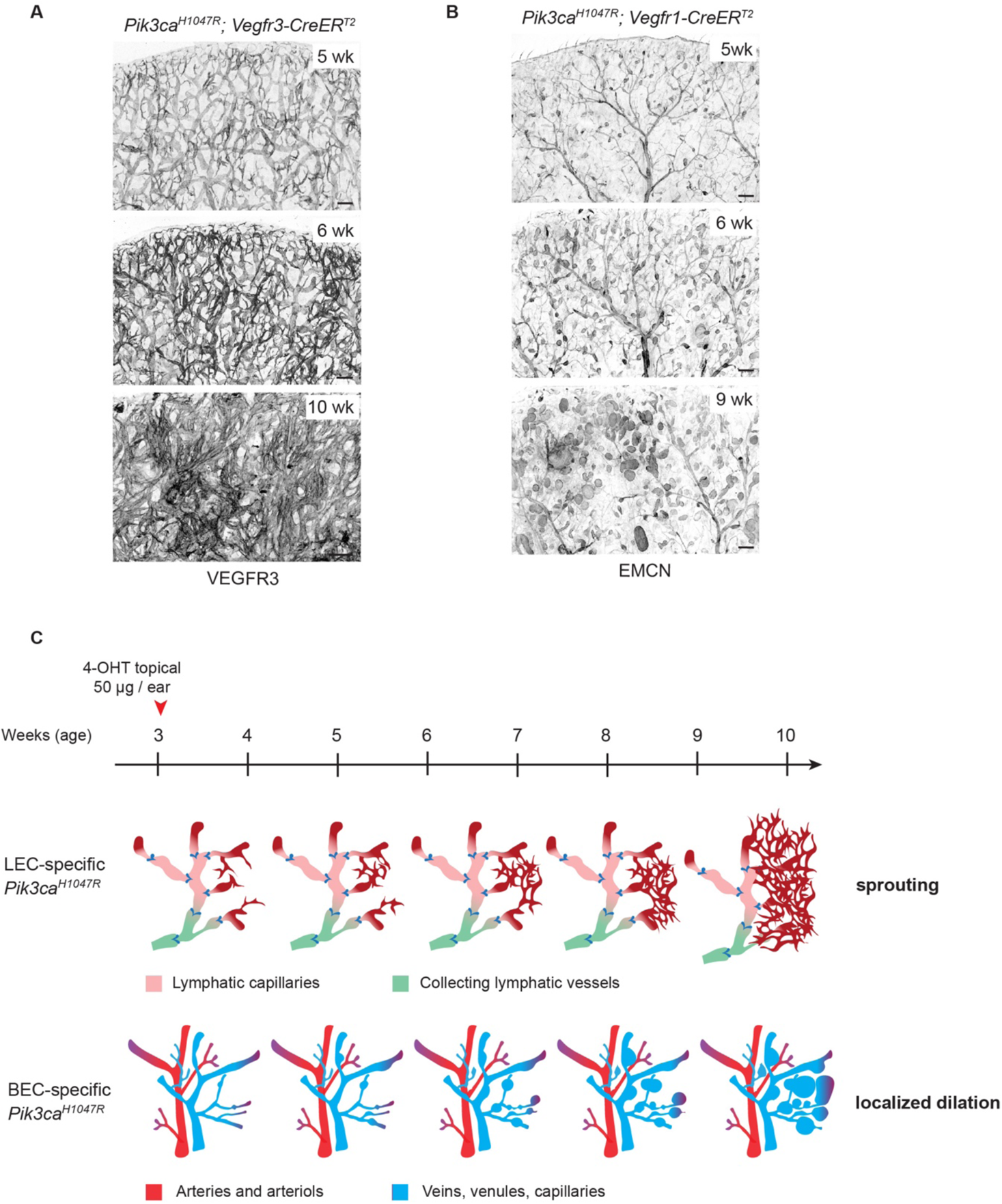
Progressive vascular overgrowth in the *Pik3ca^H1047R^; Vegfr3-CreER^T2^* and *Pik3ca^H1047R^; Vegfr1-CreER^T2^* mice. Ear tile scans of the chosen time points from time-course analysis showing vascular overgrowth phenotype in *Pik3ca^H1047R^;Vegfr3-CreER^T2^* (**A**) and *Pik3ca^H1047R^;Vegfr1-CreER^T2^*(**B**) mice. Whole-mount immunofluorescence of LECs (VEGFR3) or (venous) BECs (EMCN) in the ear skin reveals distinct responses in lymphatic (sprouting) and blood vessels (dilation). (**C**) Schematic representation of the key features of the *Pik3ca*-driven overgrowth in lymphatic and blood vessels. Scale bars: 200 μm.

**Supplemental Figure 4.**
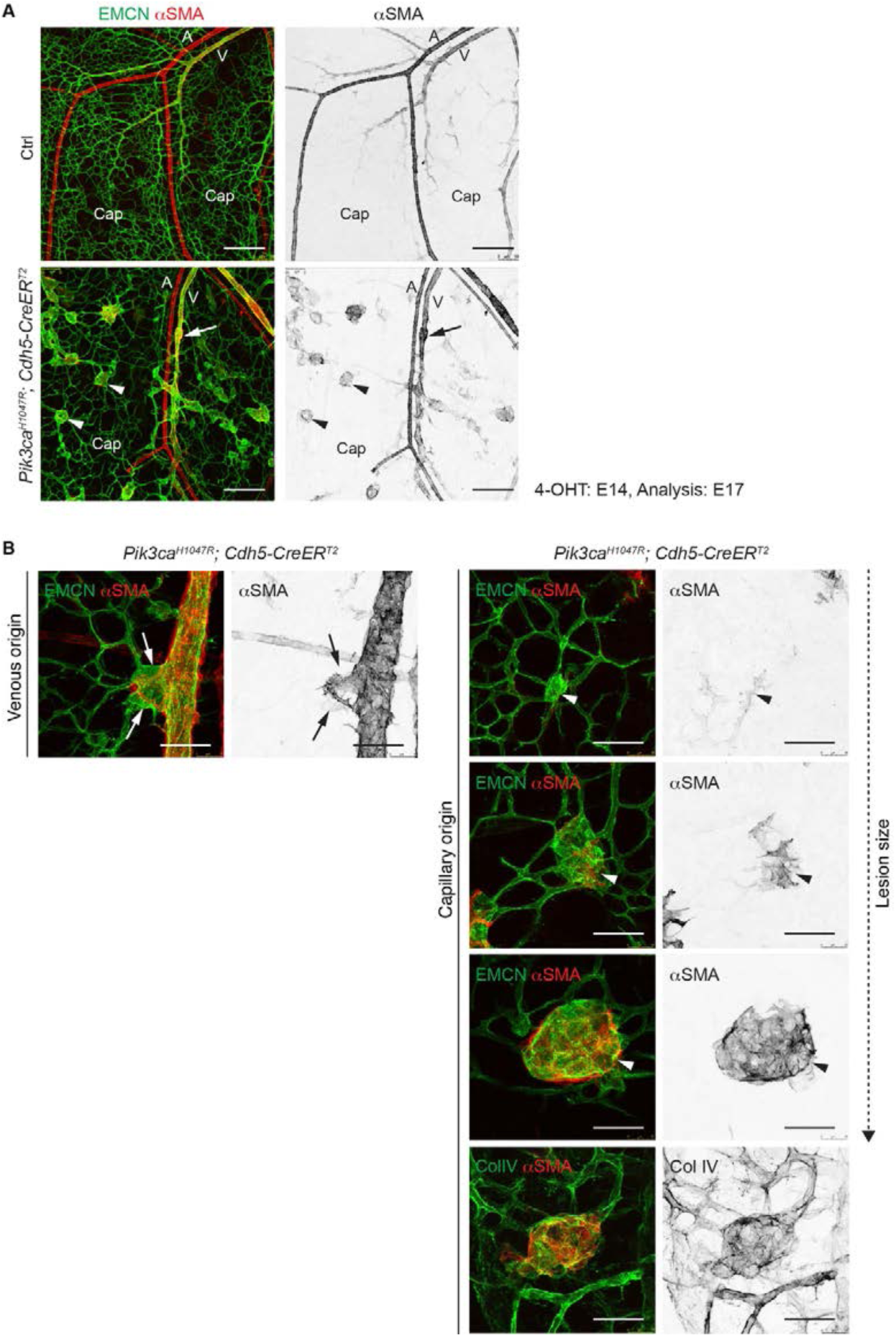
Characterization of *Pik3ca*-driven lesion formation in the embryonic blood vasculature. (**A**) Whole-mount immunofluorescence of E17 skin from *Pik3ca^H1047R^*;*Cdh5-CreER^T2^* and littermate control (Ctrl) embryos treated with 4-OHT at E14. Lesions are present in the EMCN^+^αSMA^+^ vein (V, arrow) and capillaries (Cap, arrowheads), with the latter showing ectopic αSMA coverage not present in normal capillaries. αSMA^high^ arteries (A) are not affected. (**B**) Whole-mount immunofluorescence of the skin of E15 *Pik3ca^H1047R^;Cdh5-CreER^T2^* embryos treated with 4-OHT at E11 and stained for the indicated antibodies. Single channel images are shown for the indicated stainings. Lesions originating from large veins show disruption of αSMA^+^ SMC layer (arrow) whereas lesions that originate from αSMA^−^ capillaries (arrowheads) ectopically recruit SMCs, and deposit the basement membrane protein Col IV. Scale bars: 200 µm (**A**), 50 µm (**B**).

**Supplemental Figure 5.**
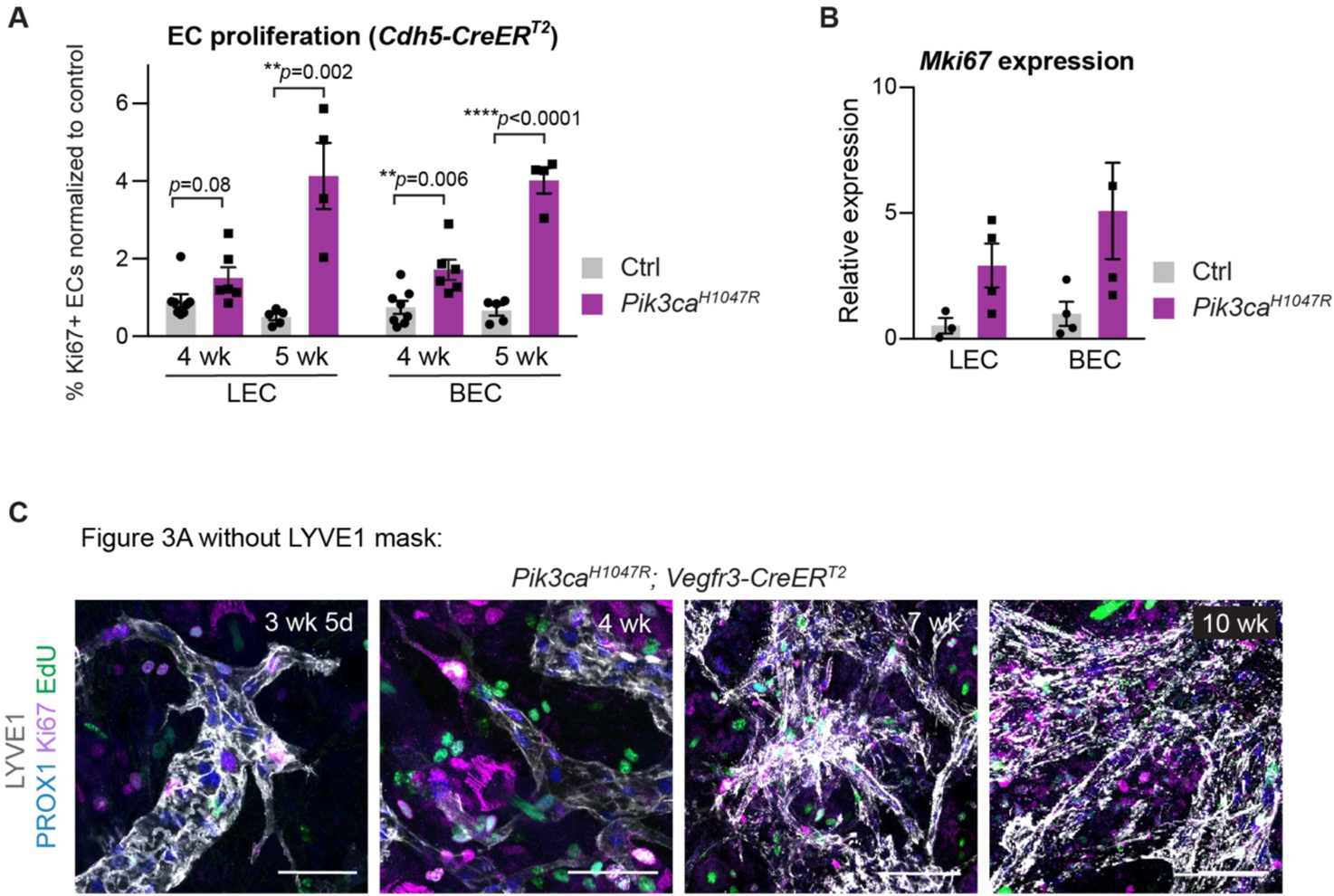
Analysis of EC proliferation during *Pik3ca*-driven vascular overgrowth. (**A**) Flow cytometry analysis of proliferating dermal LECs and BECs in *Pik3ca^H1047R^;Cdh5-CreER^T2^* and littermate control mice one or two weeks after 4-OHT treatment (corresponding 4 wk or 5 wk of age respectively). Data represent mean % of Ki67^+^ ECs, normalized to the control (*n*=4-8 mice) ± s.e.m. (**B**) qRT-PCR analysis of *Mki67* in dermal LECs and BECs, FACS-sorted from the ear skin of 4-OHT-treated 5-week-old *Pik3ca^H1047R^;Cdh5-CreER^T2^* and littermate control mice. Data represent mean relative expression (normalized to *Hprt*; *n*=3-4 mice) ± s.e.m. Transcript levels are presented relative to control BECs and LECs. (**C**) The original unmasked images for Figure 3A, showing whole-mount immunofluorescence of ear skin analyzed at different stages after 4-OHT administration in *Pik3ca^H1047R^;Vegfr3-CreER^T2^* mice. Edu was administered 16 hours prior to analysis. Scale bar: 50 μm (C).

**Supplemental Figure 6.**
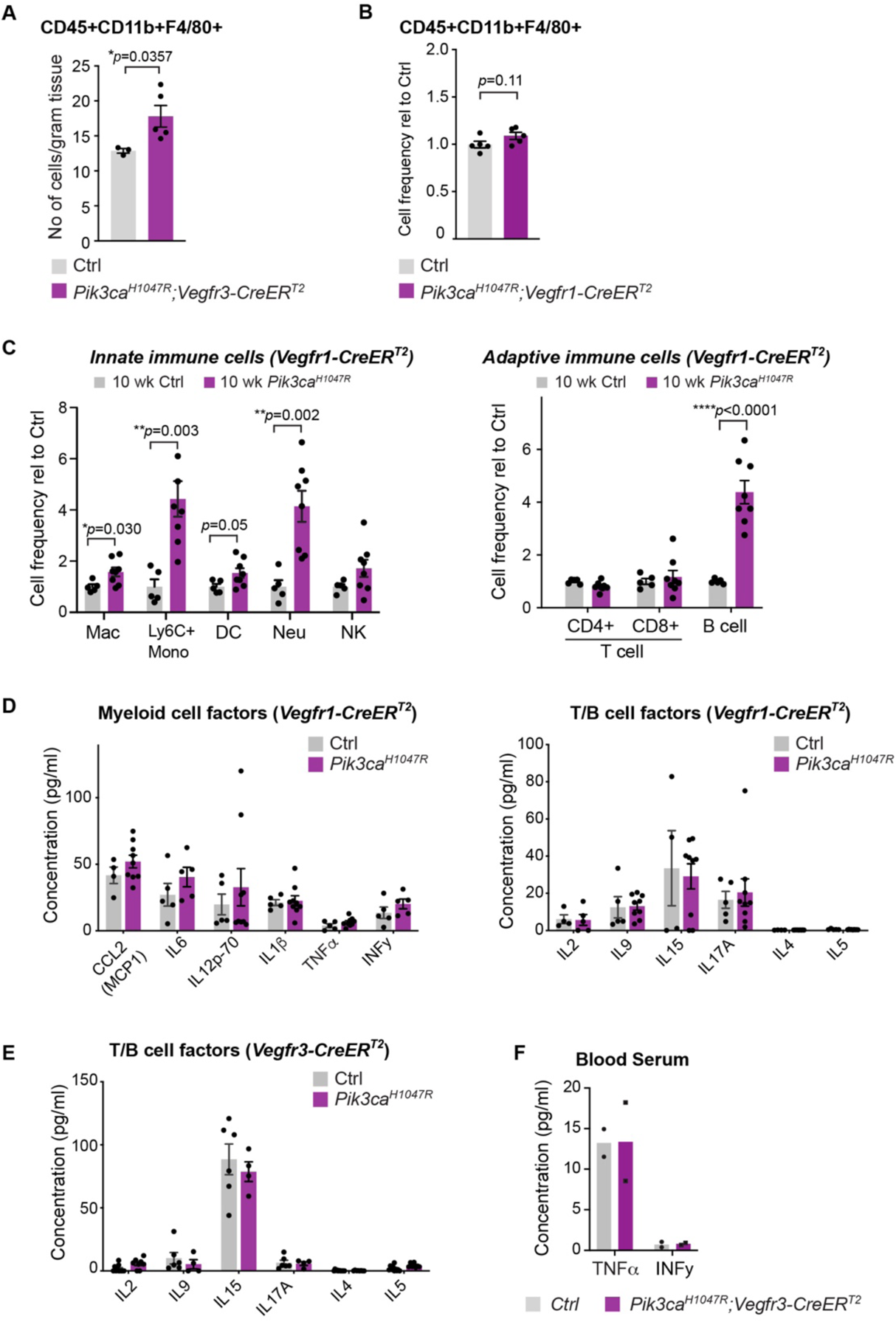
Analysis of inflammatory cells and markers in *Pik3ca*-driven vascular lesions. (**A**) Flow cytometry analysis of the number of CD45^+^CD11b^+^F4/80^+^ macrophages in the ear skin of 4-OHT-treated 5-week-old *Pik3ca^H1047R^;Vegfr3-CreER^T2^* (*n*=5) and control (*n*=3) mice. Data represent mean cell number per gram tissue ± s.e.m. *p*-value, Mann-Whitney U Test. (**B**) Flow cytometry analysis of the frequency of CD45^+^CD11b^+^F4/80^+^ antigen-presenting myeloid cells in the ear skin of 4-OHT-treated 5-week-old *Pik3ca^H1047R^;Vegfr1-CreER^T2^* (*n*=6) and control (*n*=5) mice. Data represent relative cell frequency (of live cells) relative to the control ± s.e.m. *p*-value obtained using Two-tailed unpaired Student’s t-test. (**C**) Flow cytometry analysis of innate (left) and adaptive (right) immune cells in the ear skin of 4-OHT-treated 10-week-old *Pik3ca^H1047R^; Vegfr1-CreER^T2^* mice and littermate controls. Mac, macrophage; Mono, monocyte; DC, dendritic cell; Neu, neutrophil; NK, natural killer cell. Data represent relative cell frequency (of live cells) relative to the control (*n*=5-8 mice for innate panel, n=7-9 mice for adaptive panel) ± s.e.m. *p*-value obtained using Two-tailed unpaired Student’s t-test. (**D, E**) Multiplex ELISA analysis of pro-inflammatory cytokines and chemokines associated with recruitment and/or activation of myeloid cells or T-cells and B-cells in whole ear skin lysates from *Pik3ca^H1047R^;Vegfr1-CreER^T2^* (**D**) and *Pik3ca^H1047R^;Vegfr3-CreER^T2^* (**E**) mice, and respective littermate controls. (**F**) Similar analysis of TNFα and INFγ in blood serum of *Pik3ca^H1047R^;Vegfr3-CreER^T2^* mice. Data in (**D-E**) represent mean protein levels (*n*=3-9 mice) ± s.e.m.

**Supplemental Figure 7.**
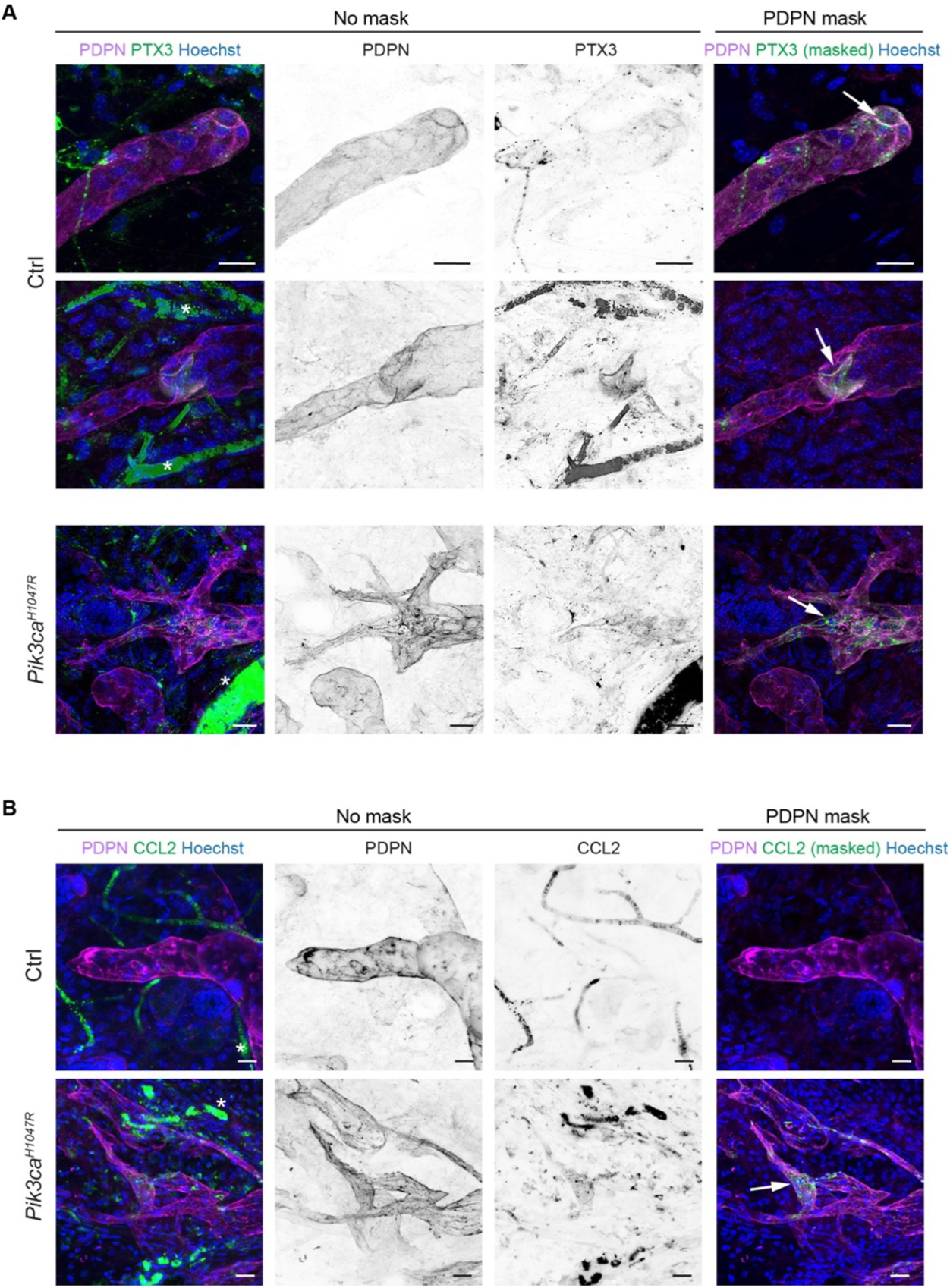
Expression of iLEC markers in normal and *Pik3ca^H1047R^*-expressing vasculature. Whole-mount immunofluorescence of control and *Pik3ca^H1047R^;Vegfr3-CreER^T2^* ear skin showing high expression of PTX3 (**A**) and CCL2 (**B**) in the capillary terminals (Ctrl) and abnormal lymphatic sprouts (mutant) (arrows). PTX3 staining is also observed in lymphatic valves (arrow). IMARIS surface mask based on PDPN expression was used to extract LEC-specific signals (images on the right, shown in the main figures). The original images including single channels for PDPN and PTX3 (A) or CCL2 (B) are shown on the left (no mask). Unspecific signal from red blood cells is indicated by asterisks. Scale bars: 20 µm.

**Supplemental Figure 8.**
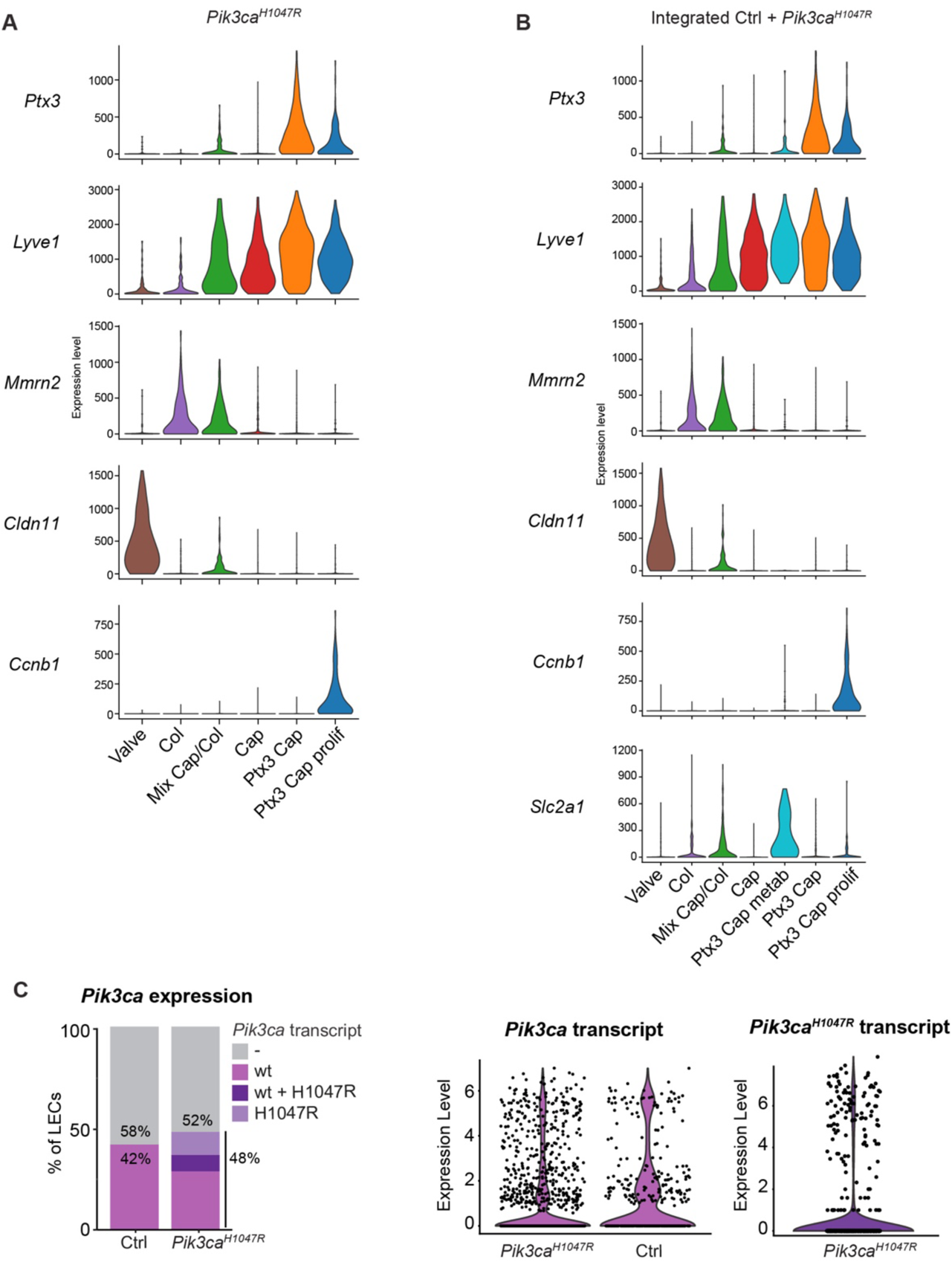
Annotation of cell clusters in scRNA-seq dataset of *Pik3ca^H1047R^* mutant LECs, and integrated control and mutant LECs. Violin plots showing the expression of selected LEC and LEC subtype marker genes in LEC clusters from the *Pik3ca^H1047R^* mutant dataset (from Figure 6A) (**A**) and integrated dataset containing LECs from control and *Pik3ca^H1047R^;Cdh5-CreER^T2^* mice (from Figure 6D) (**B**). (**C**) Proportion of LECs from control and *Pik3ca^H1047R^;Cdh5-CreER^T2^* mice expressing the endogenous mouse *Pik3ca* transcript (control and mutant mice), and/or the transgenic *Pik3ca^H1047R^* transcript (mutant mice only). Violin plots on the right show the expression levels. Note that the majority of LECs lack the *Pik3ca* transcript.

**Supplemental Figure 9.**
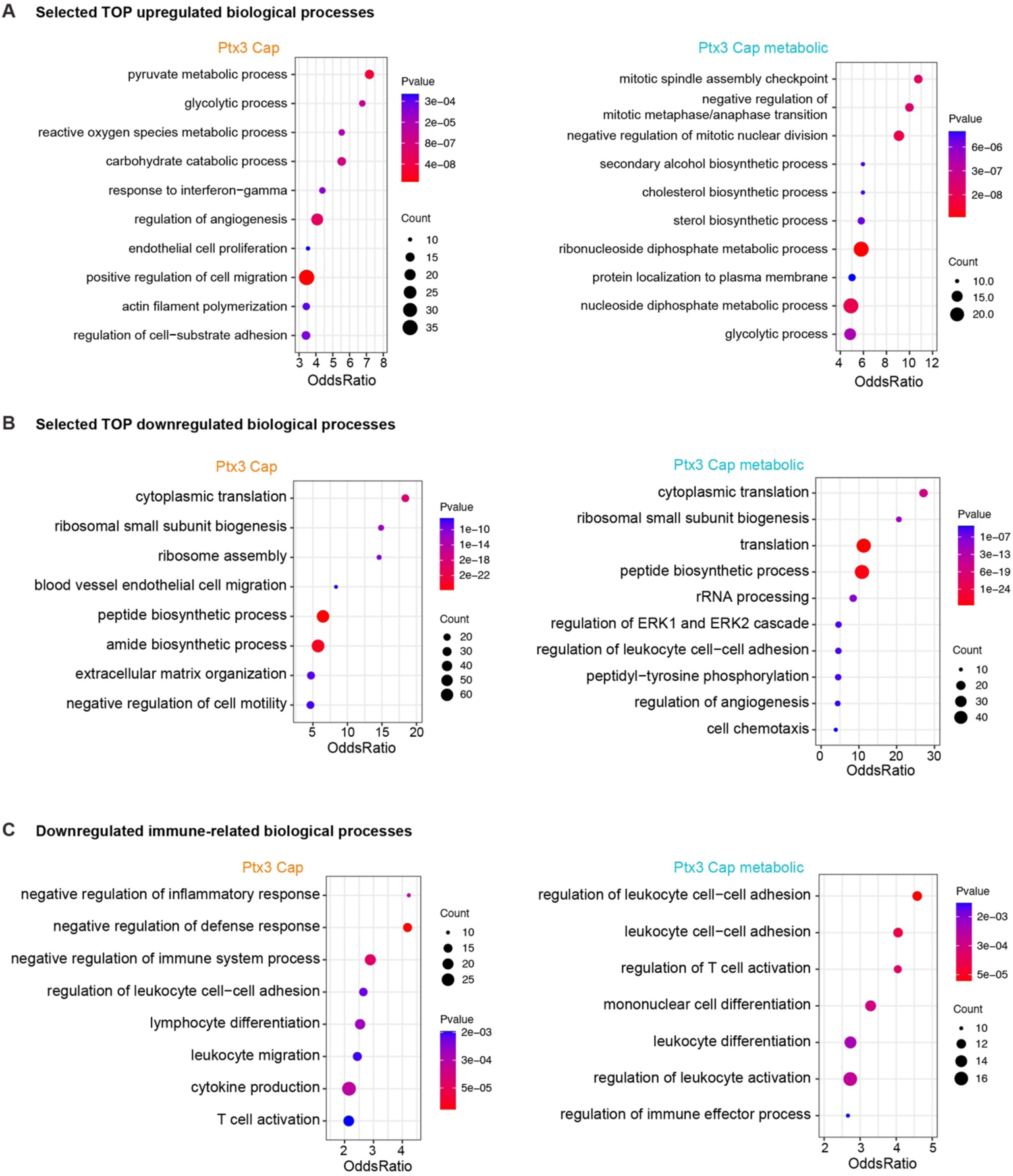
Gene Ontology enrichment analysis of differentially expressed genes in non-proliferative *Ptx3* LECs from *Pik3ca^H1047R^* mutant and control mice. Gene ontology (GO) analysis of genes upregulated (**A**) or downregulated (**B, C**) *Ptx3+* capillary LECs from *Pik3ca^H1047R^* in comparison to control mice. Selected Top terms (**A, B**) and immune-related terms (**C**) are shown Dot size illustrates count number and color illustrates adjusted p-value.

**Supplemental Figure 10.**
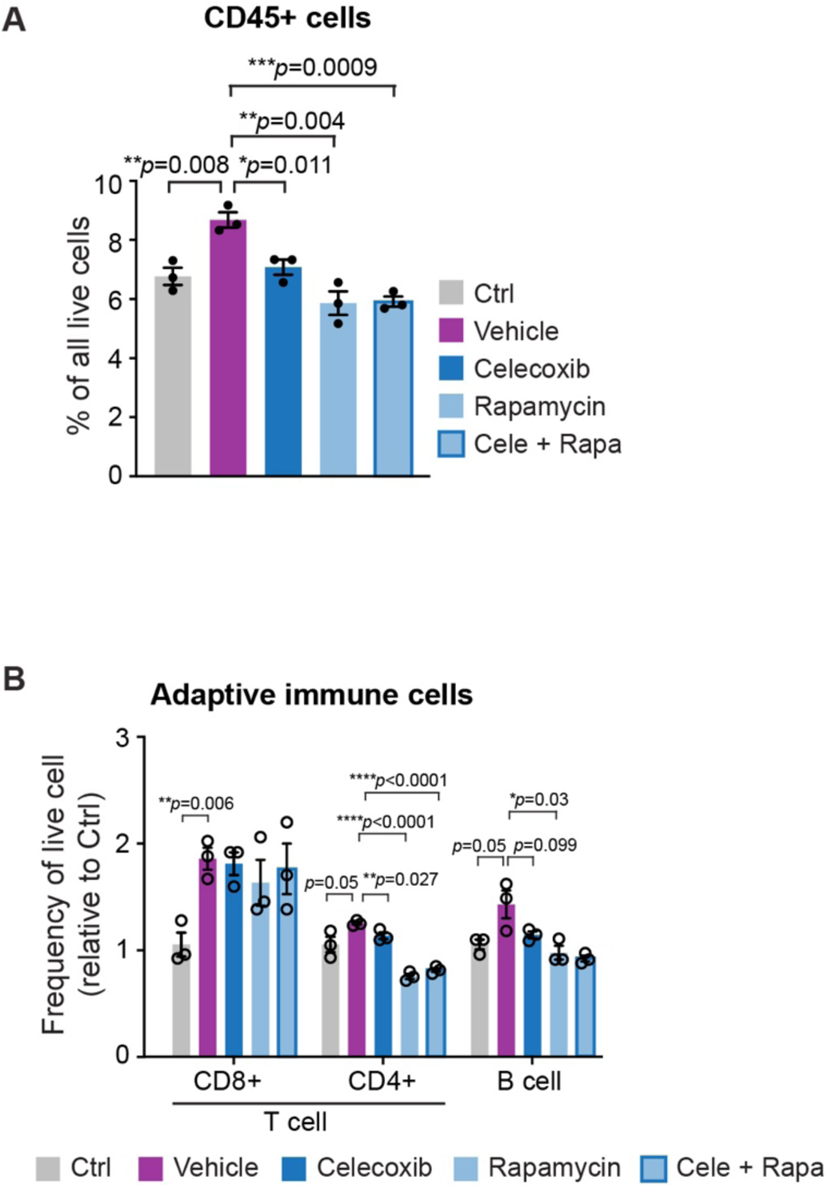
Anti-inflammatory treatment in *Pik3ca^H1047R^*-driven LM. (**A, B**) Flow cytometry analysis of CD45^+^ cells (**A**) and adaptive immune cells (**B**) in the ear skin of 4-OHT-treated 8-week-old *Pik3ca^H1047R^; Vegfr3-CreER^T2^* mice and littermate controls following treatment as in Figure 9G. Mac, macrophage; Mono, monocyte; DC, dendritic cell; Neu, neutrophil; NK, natural killer cell. Data represent relative cell frequency (of live cells) relative to the control (*n*=3 mice) ± s.e.m. *p* value, Two-tailed unpaired Student’s t-test.

**Supplemental Figure 11.**
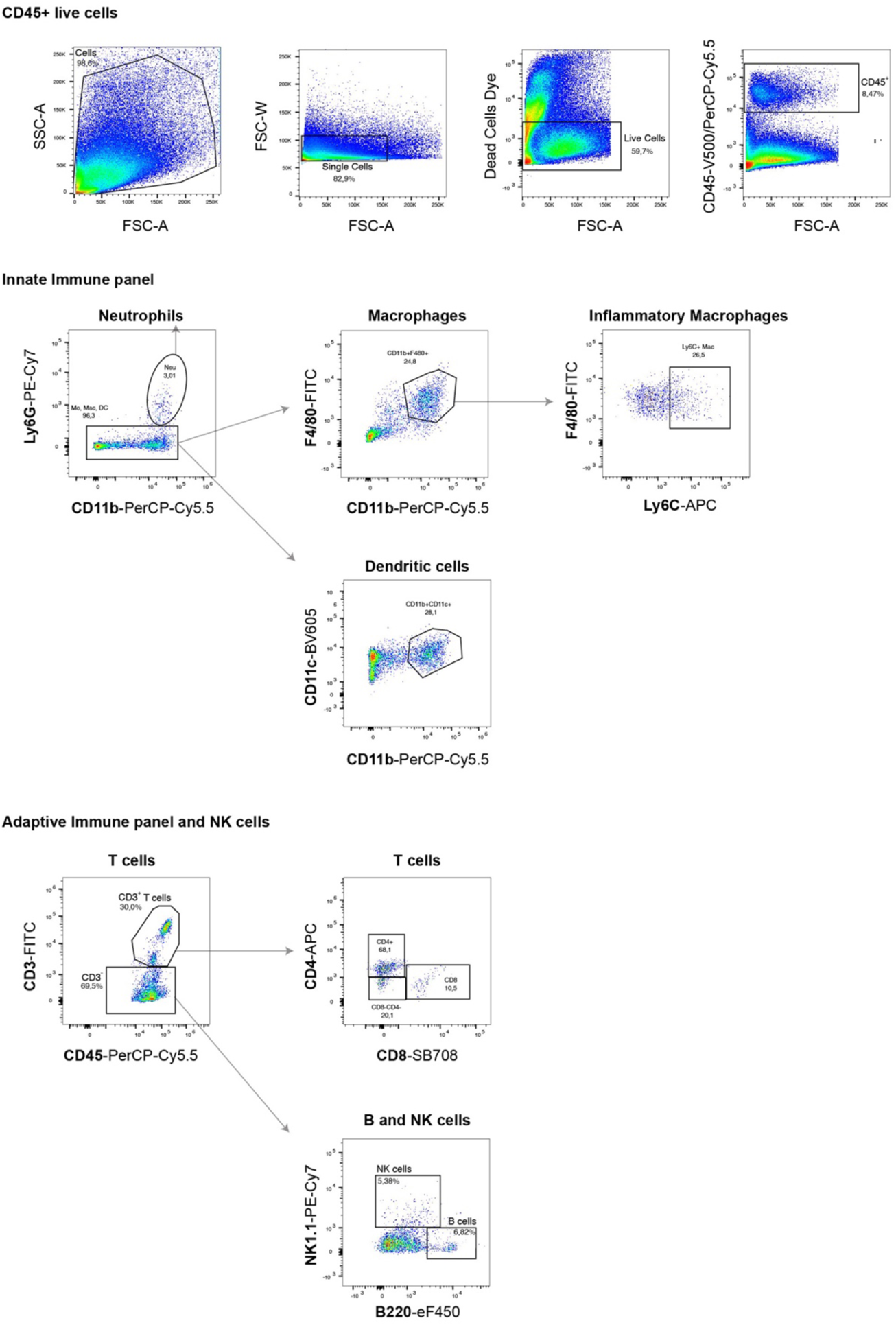
Analysis of dermal immune cell populations. Gating scheme for flow cytometry analysis of dermal innate and adaptive immune cells.

